# *RFX6* haploinsufficiency predisposes to diabetes through impaired beta cell functionality

**DOI:** 10.1101/2023.11.15.567202

**Authors:** Hazem Ibrahim, Diego Balboa, Jonna Saarimäki-Vire, Hossam Montaser, Oleg Dyachok, Per-Eric Lund, Muhmmad Omar-Hmeadi, Jouni Kvist, Om Prakash Dwivedi, Väinö Lithovius, Tom Barsby, Vikash Chandra, Solja Eurola, Jarkko Ustinov, Tiinamaija Tuomi, Päivi J. Miettinen, Sebastian Barg, Anders Tengholm, Timo Otonkoski

## Abstract

Regulatory factor X 6 (RFX6) is indispensable for pancreatic endocrine development and differentiation. The RFX6 protein-truncating variant p.His293LeufsTer7 is significantly enriched in the Finnish population with almost 1:250 individuals as a carrier. Importantly, the FinnGen study indicates a high predisposition for heterozygous carriers to develop type 2 diabetes (T2D) and gestational diabetes. To understand the role of this variant in β-cell development and function, we generated allelic series of isogenic pluripotent stem cell models and directed them into pancreatic islet lineages (SC-islets). Expectedly, *in-vitro* models of the homozygous *RFX6*^−/−^ variant failed to generate pancreatic endocrine cells, recapitulating the phenotype in Mitchell-Riley syndrome. Notably, heterozygous *RFX6*^+/−^ derived SC-islets showed reduced β-cell maturation markers and calcium oscillations, resulting in defective insulin secretion, without affecting β-cell number or insulin content. The reduced insulin secretion is sustained during *in-vivo* implantation studies, consistent with the susceptibility of the carriers to develop diabetes.

**Teaser:** Modeling *RFX6*-assocciated neonatal and type-2 diabetes using allelic series stem cell-derived islets *in-vitro* and *in-vivo*.

## Introduction

Diabetes is a metabolic disorder characterized by the inability of pancreatic β-cells to control blood glucose levels. The etiology of the most common form, type 2 diabetes (T2D), is heterogeneous, although population genetic studies have identified numerous T2D-associated genetic variants in loci of genes expressed in the β-cell (*1*, *2*). Pathogenic variants in these β-cell genes lead to monogenic diabetes, presenting either as neonatal diabetes or maturity onset diabetes of the young (MODY) (*3*). Regulatory factor X 6 (RFX6) is a winged-helix transcription factor that regulates genes required for the development of the pancreas and other intestinal organs (*4–7*). Autosomal recessive variants in *RFX6* cause Mitchell-Riley syndrome (OMIM, 615710), characterized by intrauterine growth retardation, annular or hypoplastic pancreas, permanent neonatal diabetes, gall bladder hypoplasia or agenesis, intestinal stenosis with malabsorptive diarrhea and in some cases, pancreatic exocrine insufficiency (*6*, *8*, *9*). A milder form of Mitchell-Riley syndrome was recently reported with compound heterozygous variants that are not completely inactivating, resulting in a later-onset diabetes (between the age of 2–5 years) (*10*, *11*).

In mice, Rfx6 is initially expressed in the definitive endoderm, before becoming progressively confined to the gut and dorsal pancreatic bud, and then islet progenitor cells where it is required for the differentiation of all islet cell types, except for pancreatic polypeptide-producing PP cells. Loss of Rfx6 in mice showed similarities to the human phenotype, including neonatal diabetes and intestinal obstruction, but with variable pancreatic hypoplasia (*6*). While homozygous Rfx6 mutant mice had severe symptoms and died shortly after birth, heterozygous mutants did not show any signs of diabetes (*6*). Conditional Rfx6 depletion in adult mouse β-cells showed impaired glucose-stimulated insulin secretion (GSIS), without compromising β-cell mass or insulin content. The defective insulin secretion was attributed to the downregulation of β-cell maturation genes *Gck*, *Abcc8*, *Ucn3* and voltage-dependent calcium channels (VDCCs), concomitant with the upregulation of β-cell disallowed genes such as *Slc16a1*, *Ldha*, and *Igfbp4* (*12*). Similarly, knockdown of *RFX6* in the human β-cell insulinoma line EndoC-βH2 resulted in reduced insulin gene transcription and defective glucose stimulated insulin secretion through reducing VDCC gene expression (*13*). A recent study also demonstrated that *RFX6* knockdown in primary human islets reduced GSIS to the level seen in T2D islets, through transcriptionally dysregulated vesicle trafficking, exocytosis, and ion transport pathways (*14*).

Heterozygous *RFX6* pathogenic variants have been linked to MODY with reduced penetrance in humans (*15–21*). Moreover, genome-wide association studies (GWAS) have associated variants of *RFX6* with T2D (*22*, *23*). In adult primary islets, RFX6 was shown to be a hub transcription factor that was downregulated in early T2D islets, which was correlated with reduced GSIS (*14*). Additionally, genetic variants that increase T2D risk are predicted to disrupt RFX-binding motifs (*24*).

The precise mechanism of how heterozygous pathogenic *RFX6* variant carriers are predisposed to develop diabetes remains unknown. Therefore, we sought to use patient- and embryonic-stem cells combined with CRISPR-based genetic engineering to create isogenic allelic series models of a specific *RFX6* protein-truncating variant (PTV); circumventing the use of unphysiological systems of complete gene knockout or knockdown. Human pluripotent stem cells have been extensively used to model monogenic diabetes, as they can be differentiated into stem-cell-derived islets (SC-islets) that closely mimic native human islets developmentally and functionally (*25–27*). Utilizing our state-of-the-art protocol to generate highly functional SC-islets (*28*, *29*), we show that homozygous *RFX6* PTV leads to sustained SOX9 and NEUROG3 expression leading to increased apoptosis and deviating the endocrine lineage specification of pancreatic endocrine cells. Importantly, we demonstrate how *RFX6* haploinsufficiency in heterozygous SC-islets leads to impaired insulin secretion capacity *in-vitro* and *in-vivo*, through reducing β-cell maturation markers and intracellular Ca^2+^ levels.

## Results

### Impact of heterozygous and homozygous *RFX6* PTV p.His293LeufsTer7 on diabetes development

A two-nucleotide deletion in exon 9 (c.878_879delAC) of the *RFX6* gene leads to a frame shift at codon 293 and an early stop codon at 298 (p.His293LeufsTer7), instead of the 928 amino acid wild-type protein (*30*). While the wild-type protein contains a winged-helix RFX DNA-binding domain and three dimerization domains mediating both hetero- and homo-dimeric interactions (*6*), this protein-truncating variant (PTV) is predicted to contain only the DNA binding domain and the first dimerization domain (Fig. 1A). We assessed the impact of this heterozygous PTV on the lifetime risk of T2D and gestational diabetes in the FinnGen dataset (data freeze R11), containing data from 440,734 individuals. It included 1,318 heterozygous carriers of the variant, who showed 80% higher risk of T2D (p=6.8×10^−28^) (with on average a 2-year earlier age of onset) (Fig. 1B) and gestational diabetes (79%, p=2.3×10^−8^) (Fig. 1C). Interestingly, the carriers also demonstrated a reduced body mass index (Beta=-0.12 SD, p=1.3×10^−4^ and n=321,672) compared to non-carriers.

**Fig. 1.**
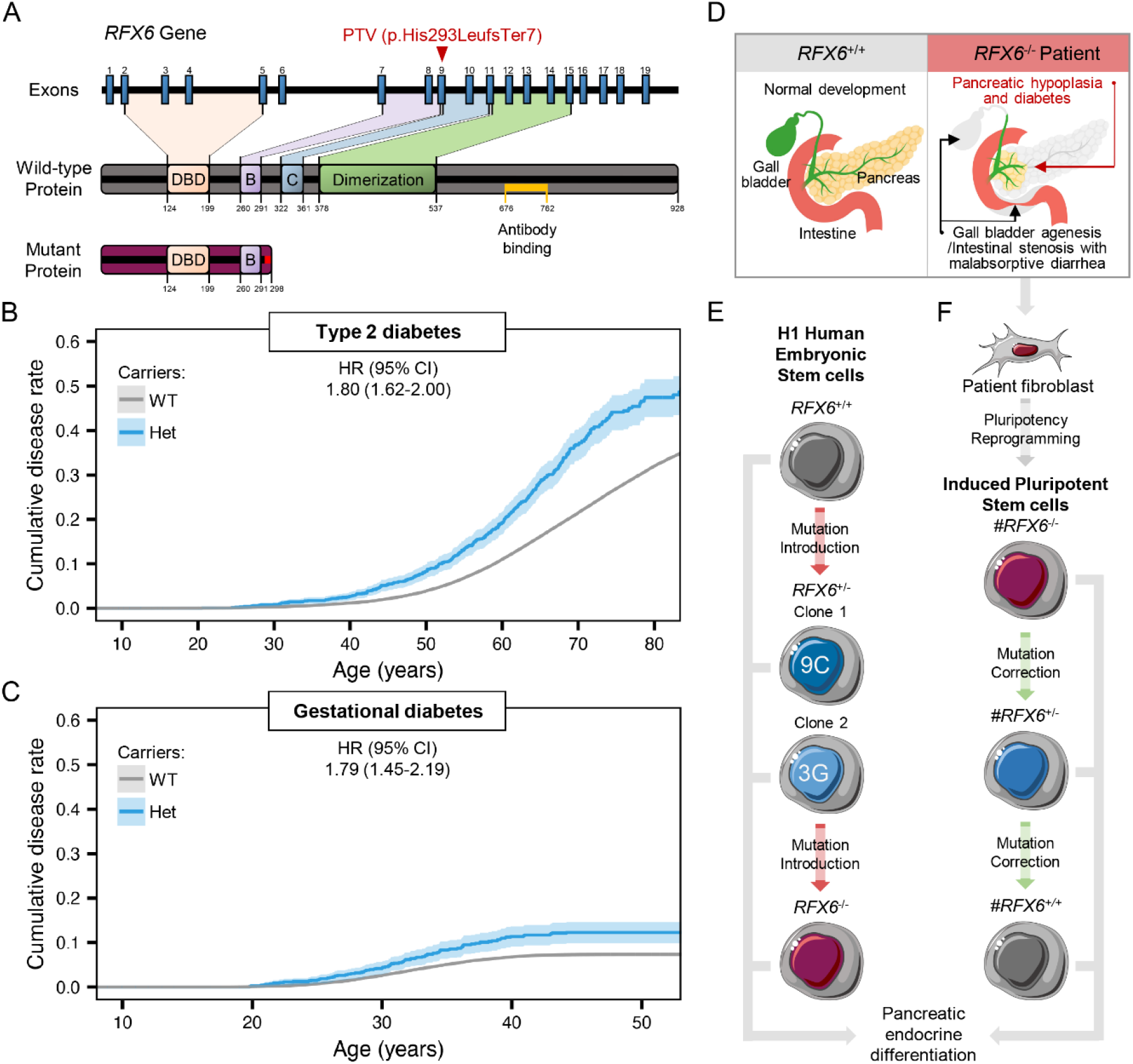
Impact of heterozygous and homozygous *RFX6* PTV p.His293LeufsTer7 on diabetes development in the Finnish population. **(A)** The human *RFX6* gene and the encoded wild-type (WT) and protein-truncating variant (PTV) proteins. The WT RFX6 is a 928-amino acid protein containing a DNA-binding domain (DBD) and three dimerization domains (B, C and Dimerization). The RFX6 PTV p.His293LeufsTer7 contains the DBD, dimerization domain B and a frameshift making an early stop codon at 298. The antibody used in this study binds to the amino acid sequence from 676-762 (shown in yellow). **(B)** Survival curve stratified by *RFX6* genotype (WT; wild-type 439,416 and Het; heterozygous carriers 1,318) and adjusted hazard ratio (HR) for type 2 diabetes risk. The mean (SD) age at onset for mutation carriers (Het) and non-carriers (WT) was 57.48 (12.87) and 59.91 (12.43) years, respectively (p-value=2.5×10^−4^). The total number of individuals was 440,734 which included 71,728 with type 2 diabetes. **(C)** Survival curve stratified by *RFX6* genotype (WT; wild-type, n=253,803 and Het; heterozygous carriers, n=814) and adjusted hazard ratio (HR) for gestational diabetes risk. The total number of individuals was 254,617 which included 16,802 with gestational diabetes. **(D)** Schematic showing the patient’s clinical manifestations (pancreatic hypoplasia with neonatal diabetes, gall bladder agenesis and intestinal stenosis with malabsorptive diarrhea). **(E)** Schematic of generating *RFX6* allelic series of the mutation in H1 human embryonic stem cells (hESCs), followed by directed differentiation into pancreatic endocrine cells. **(F)** Schematic of patient derived fibroblasts reprogramming to induced pluripotent stem cells (iPSCs), followed by generating *RFX6* allelic series of the mutation, and directed differentiation into pancreatic endocrine cells. Survival plots in (B and C) were generated using survminer and adjusted hazards ratios were calculated using Cox proportional hazards model adjusting for age, sex and PC1-PC10.

We identified one individual carrying this PTV in homozygous form. This male patient is currently alive at 9 years of age. He was diagnosed with Mitchell-Riley syndrome at birth, manifesting pancreatic hypoplasia, neonatal diabetes, gall bladder agenesis, intestinal stenosis and malabsorptive diarrhea (Fig. 1D). The patient had low birth size, which recovered gradually after administering total parenteral nutrition (Supplementary Fig. 1A). At the age of 4 years, magnetic resonance cholangiopancreatography showed a hypoplastic pancreas of 4 mL (data not shown); the average volume is 20 mL for healthy individuals of the same age (*31*). Blood glucose was normal at birth, but became hyperglycemic at the age of 4 days. Insulin treatment was started with an insulin dose of 0.1 IU/kg/day, since C-peptide has been barely detectable (Supplementary Fig. 1B). Plasma glucagon was undetectable at birth, but curiously, it normalized gradually over the course of 4 months (Supplementary Fig. 1C). In contrast, pancreatic polypeptide dropped below the detection limit (Supplementary Fig. 1D). Pancreatic exocrine function has been severely impaired (Supplementary Fig. 1E). Since the clinical features of this homozygous PTV represented typical Mitchell-Riley syndrome, we considered it as a loss-of-function variant *RFX6*^−/−^.

To study this PTV in more detail, we generated two isogenic allelic series stem cell models. In the first model, we introduced the PTV in H1 human embryonic stem cells (hESCs). We validated two heterozygous clones (*RFX6*^+/−^ 9C and *RFX6*^+/−^ 3G) and one homozygous (*RFX6*^−/−^ 1H) (Fig. 1E and Supplementary Fig. 2). In the second model, we generated patient-derived induced pluripotent stem cells (iPSCs) from the homozygous patient and then corrected the PTV heterozygously and homozygously, creating the iPSC-lines (#*RFX6*^−/−^, *#RFX6*^+/−^ and *#RFX6*^+/+^) (Fig. 1F and Supplementary Fig. 3).

### RFX6 controls the transcriptional network of pancreatic development

We differentiated the generated hESCs towards functional SC-islets using our optimized 7-stage protocol (*28*, *29*) (Fig. 2A). Since *RFX6* expression started to increase at the posterior foregut stage (S3) (Fig. 2B), the PTV did not affect the earlier stage of definitive endoderm induction (data not shown). However, levels of *RFX6* gene expression in both heterozygous *RFX6*^+/−^ and homozygous *RFX6*^−/−^ cells were reduced (Fig. 2B). The loss of RFX6 in *RFX6*^−/−^ cells was confirmed by immunoblotting at S3, and interestingly, both heterozygous clones showed significant reduction in RFX6 protein levels at this stage, indicating haploinsufficiency (Fig. 2C, D). The lack of RFX6 in *RFX6*^−/−^ cells led to a significant reduction in *PDX1* expression, a key transcription factor in pancreatic development (Fig. 2E). Subsequently, the percentage of PDX1^+^ NKX6.1^+^ and the emergence of CHGA^+^ cells were significantly reduced (all markers of pancreatic endocrine lineages), while both heterozygous clones were similar to *RFX6*^+/+^ (Fig. 2F and Supplementary Fig. 4). Since the quantity and quality of pancreatic progenitors and endocrine precursors were impaired in *RFX6*^−/−^, but not in *RFX6*^+/−^, we ran bulk RNA sequencing analysis (RNAseq) to identify differentially expressed genes between *RFX6*^−/−^ and *RFX6*^+/+^ at stages 3 (S3) and 5 (S5) (Supplementary Fig. 5).

**Fig. 2.**
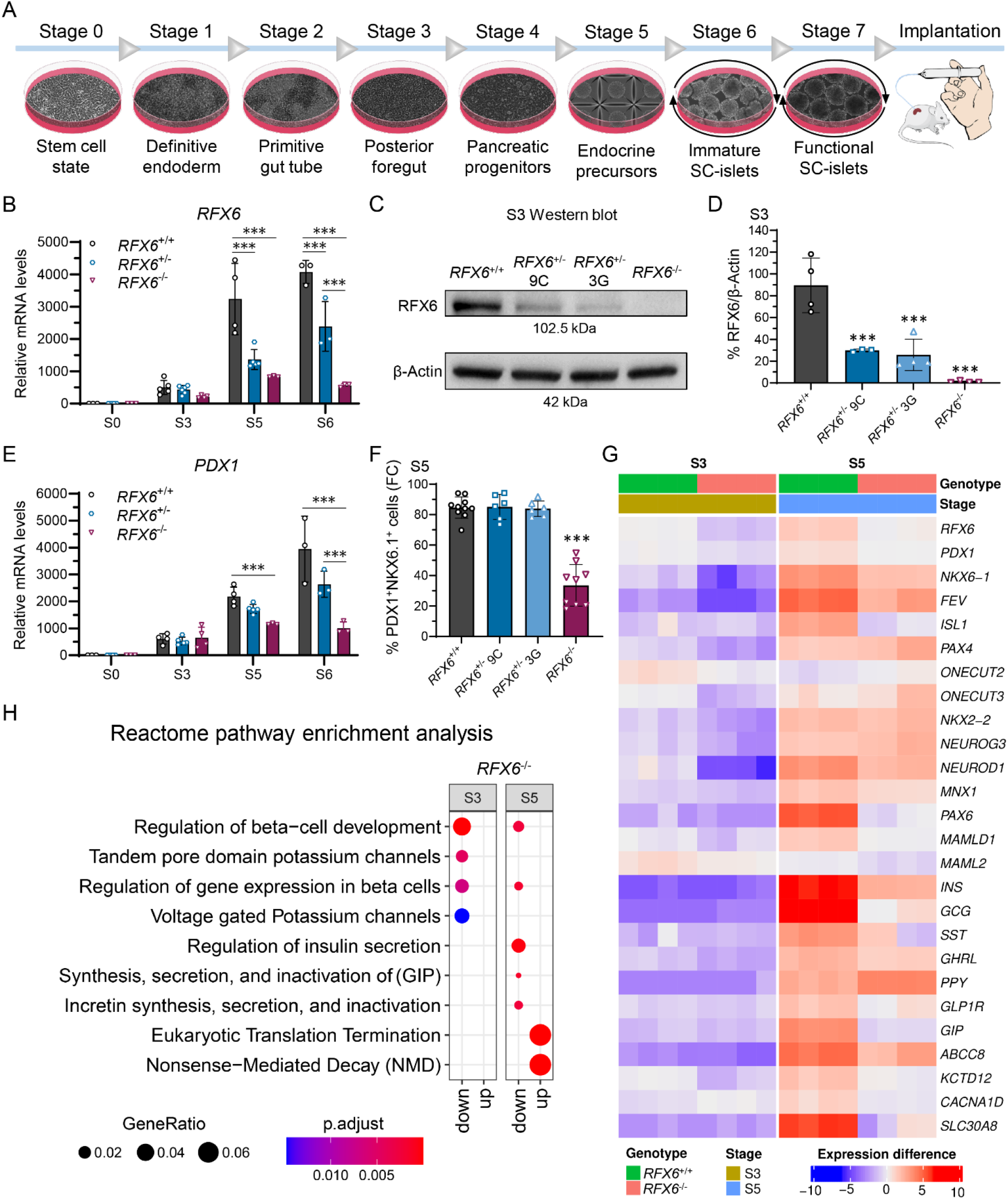
RFX6 controls the transcriptional network of pancreatic development. **(A)** Schematic of H1 SC-islet differentiation protocol. Stages 1–4 in monolayer, Stage 5 in microwells and Stages 6–7 in suspension culture, followed by implantation under kidney capsule of NSG mice. **(B)** Relative gene expression levels of *RFX6* at S0, S3, S5 and S6 (n=3–5). **(C)** Protein immunoblot for RFX6 and β-Actin for the isogenic H1 clones at S3. **(D)** Percentage of RFX6 protein bands densitometry normalized to β-Actin bands, quantified from (E) (n=3–4). **(E)** Relative gene expression levels of *PDX1* at S0, S3, S5 and S6 (n=3–5). **(F)** Percentage of PDX1^+^ and NKX6.1^+^ cells at S5 measured by flow cytometry (n=6–10). **(G)** Heatmap for bulk RNAseq showing differentially expressed genes between *RFX6^+/+^* and *RFX6^−/−^* at S3 and S5 (n=4). Each gene is shown with a multiple testing corrected p value generated for the longitudinal differential expression of the gene during differentiation. (**F**) Reactome pathway enrichment analysis between *RFX6^+/+^ and RFX6^−/−^* at S3 and S5. Statistical significance was measured using two-way ANOVA with Tukey’s test for multiple comparisons correction in (B and C), and one-way ANOVA with Tukey’s test for multiple comparisons correction in (D and F). Error bars represent ±SD from the mean, *p<0.05, **p<0.01, ***p<0.001.

In *RFX6*^−/−^ cells, the transcription factor regulatory network orchestrating the development of the pancreatic lineage was downregulated at S3 including *RFX6*, *PDX1*, *NKX6.1*, *FEV*, *PAX4*, *NEUROG3* and *NEUROD1* (Fig. 2G). While most of these genes remained downregulated at S5 in addition to *ISL1* and *PAX6*, gene expression levels of *NEUROG3*, *NKX2.2* and *PAX4* recovered and even nominally increased compared to *RFX6*^+/+^. Interestingly, all pancreatic hormone levels were downregulated at S5, except for *PPY* expression, which was significantly upregulated (Fig. 2G). In agreement with these findings, Reactome enrichment analysis showed downregulation of pathways involved in β-cell development and gene expression in *RFX6*^−/−^ cells. Additionally, voltage gated potassium channels were reduced at S3, and the regulation of insulin secretion, and GIP synthesis and secretion were reduced at S5 (Fig. 2H).

We further confirmed these findings in our isogenic iPSC model using a simplified monolayer differentiation protocol. Similar to the hESCs, the emergence of PDX1^+^ cells in #*RFX6*^−/−^ was not compromised, while *RFX6* and *PDX1* gene expression, and the generation of NKX6.1^+^ pancreatic progenitors were significantly reduced (Supplementary Fig. 6 and Supplementary Fig. 7A, B). We then investigated if the PTV-harbouring cells have reduced *RFX6* expression due to nonsense-mediated decay (NMD) of *RFX6* mRNA. We treated #*RFX6*^+/−^ cells with the NMD inhibitor cycloheximide at S3, and quantified the reads of variant and corrected alleles. In the non-treated samples, the percentage of PTV cDNA was only 15%, while treatment with cycloheximide normalized the expression of the variant allele gradually to ≈48% in a dose-dependent manner, confirming that the PTV mRNA is subjected to NMD (Supplementary Fig. 7C). At the endocrine stage (S6), #*RFX6*^−/−^ cells lacked the expression of the five pancreatic hormones, insulin, glucagon, somatostatin, pancreatic-polypeptide and ghrelin, while this phenotype was rescued in both the heterozygous and homozygous corrected cells (Supplementary Fig. 7D).

### Sustained expression of SOX9 and NEUROG3 in homozygous *RFX6*^−/−^ cells

We further studied the endocrine precursors in the 3D model of hESCs and found a peculiar dysregulation of SOX9 and NEUROG3. Normally, when SOX9 is highly expressed in the pancreatic epithelium it induces the proendocrine transcription factor NEUROG3, which upon reduction of notch signaling promotes the endocrine lineage differentiation program (*32*). Indeed, in our S4 pancreatic progenitors, *SOX9* was highly expressed across all lines (Supplementary Fig. 4A and Fig. 3B). Upregulation of NEUROG3 expression at S5 was associated with downregulation of *SOX9* expression (Fig. 3A-C). A further decrease of *SOX9* expression was noted in the immature S6 SC-islets in *RFX6*^+/+^ and *RFX6*^+/−^, but not in *RFX6*^−/−^ (Fig. 3B). Interestingly, *NEUROG3* expression nominally increased at S5 in *RFX6*^−/−^, similar to the observation in the bulk RNAseq, and was significantly higher than in the other genotypes at S6 (Fig. 3C). The pattern of SOX9 and NEUROG3 expression was similar between the genotypes in S5 endocrine precursors as shown by immunohistochemicry (Fig. 3A). However, the pattern was completely altered at S6. Whereas SOX9 was detected mainly in the cytoplasm of *RFX6*^+/+^ and *RFX6*^+/−^ cells, nuclear SOX9 expression was sustained in *RFX6*^−/−^ (Fig. 3D). The numbers of nuclear SOX9^+^ cells and NEUROG3^+^ cells were significantly increased in *RFX6*^−/−^ (Fig. 3E). Since NEUROG3 expression induces the endocrine lineage specification, we investigated the expression of the endocrine marker CHGA. While ≈80% of the cells were CHGA^+^ in *RFX6*^+/+^ and *RFX6*^+/−^, only ≈40% were detected in *RFX6*^−/−^, none of which were insulin positive (Fig. 3F). At the final stage S7, *RFX6*^−/−^ cells did not thrive for more than two days. TUNEL staining at the end of S6 revealed significantly increased apoptosis in *RFX6*^−/−^ (Fig. 3G, H), suggesting that failure to specify the pancreatic endocrine precursors due to the lack of RFX6 leads to progressive cell death.

**Fig. 3.**
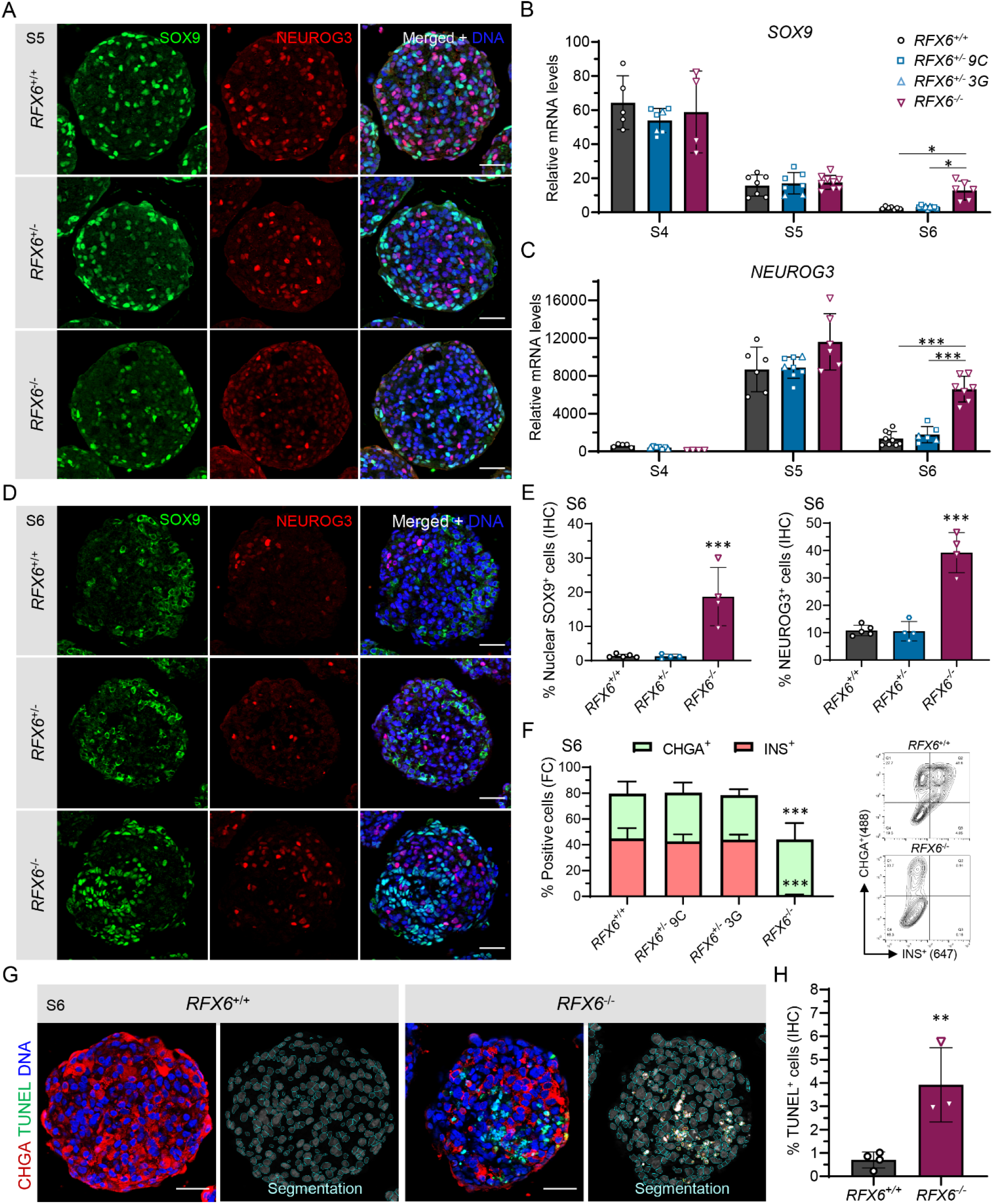
Sustained expression of SOX9 and NEUROG3 in homozygous *RFX6^−/−^* cells. **(A)** Immunohistochemistry showing SOX9^+^ and NEUROG3^+^ cells at S5, scale bars, 50 µm. **(B)** Relative gene expression levels of *SOX9* comparing all the cell-lines at S4, S5 and S6 (n=4– 9). **(C)** Relative gene expression levels of *NEUROG3* comparing all the cell-lines at S4, S5 and S6 (n=4–9). **(D)** Immunohistochemistry for SOX9^+^ and NEUROG3^+^ cells at S6, scale bars, 50 µm. **(E)** Percentages of SOX9^+^ and NEUROG3^+^ cells at S6 quantified from (D) (n=4–5). **(F)** Percentage of CHGA^+^ cells (light green) and INS^+^ cells (light red) at S6 quantified by flow cytometry (n=5–6). **(G)** Immunohistochemistry showing TUNEL^+^ and CHGA^+^ cells for the *RFX6^+/+^ and RFX6^−/−^* clones at S6, scale bars, 50 µm. **(H)** Percentage of TUNEL^+^ cells at S6 quantified from (H) (n=3–4). Statistical significance was measured using two-way ANOVA with Tukey’s test for multiple comparisons correction in (B and C), one-way ANOVA with Tukey’s test for multiple comparisons correction in (E and G), and using two-tailed unpaired t-test in (I). Error bars represent ±SD from the mean, *p<0.05, **p<0.01, ***p<0.001.

### *RFX6* haploinsufficiency impairs insulin secretion of β-cells

Since *RFX6*^−/−^ cells failed to survive past S6, we continued to study the functionality of only the WT and heterozygous cells at the mature stage S7. The proportions of INS^+^ and GCG^+^ cells (markers of beta and alpha cells respectively), and total insulin content were similar between the two genotypes (Fig. 4A-C, Supplementary Fig. 8A). We confirmed the sustained *RFX6* haploinsufficiency at this stage using protein immunoblotting (Fig. 4D, E). Glucose-stimulated insulin secretion (GSIS) in a static assay, normalized to DNA content, showed impaired insulin secretion in *RFX6*^+/−^ compared to the wild type in both basal and high glucose conditions, while KCl-stimulated insulin secretion showed no difference (Fig. 4F). Glucose stimulatory index was similar between *RFX6*^+/+^ and *RFX6*^+/−^ (Supplementary Fig. 8B). Correspondingly, dynamic perifusion assays confirmed the reduced insulin secretion in both glucose conditions. It also revealed a blunted first-phase response, and lower insulin secretion when stimulated with the GLP-1 analogue exendin-4 (Fig. 4G). Insulin secretion shut down effectively when the glucose concentration was decreased back to 2.8 mM in both genotypes, followed by a normal maximum secretion capacity with KCl depolarization. Total area under the curve (AUC) for the whole perifusion assay was significantly reduced in *RFX6*^+/−^ (Fig. 4H). This highlights that lower RFX6 expression in the heterozygous cells reduces the insulin secretion capacity in basal and stimulatory glucose conditions without affecting β-cell number or insulin content.

**Fig. 4.**
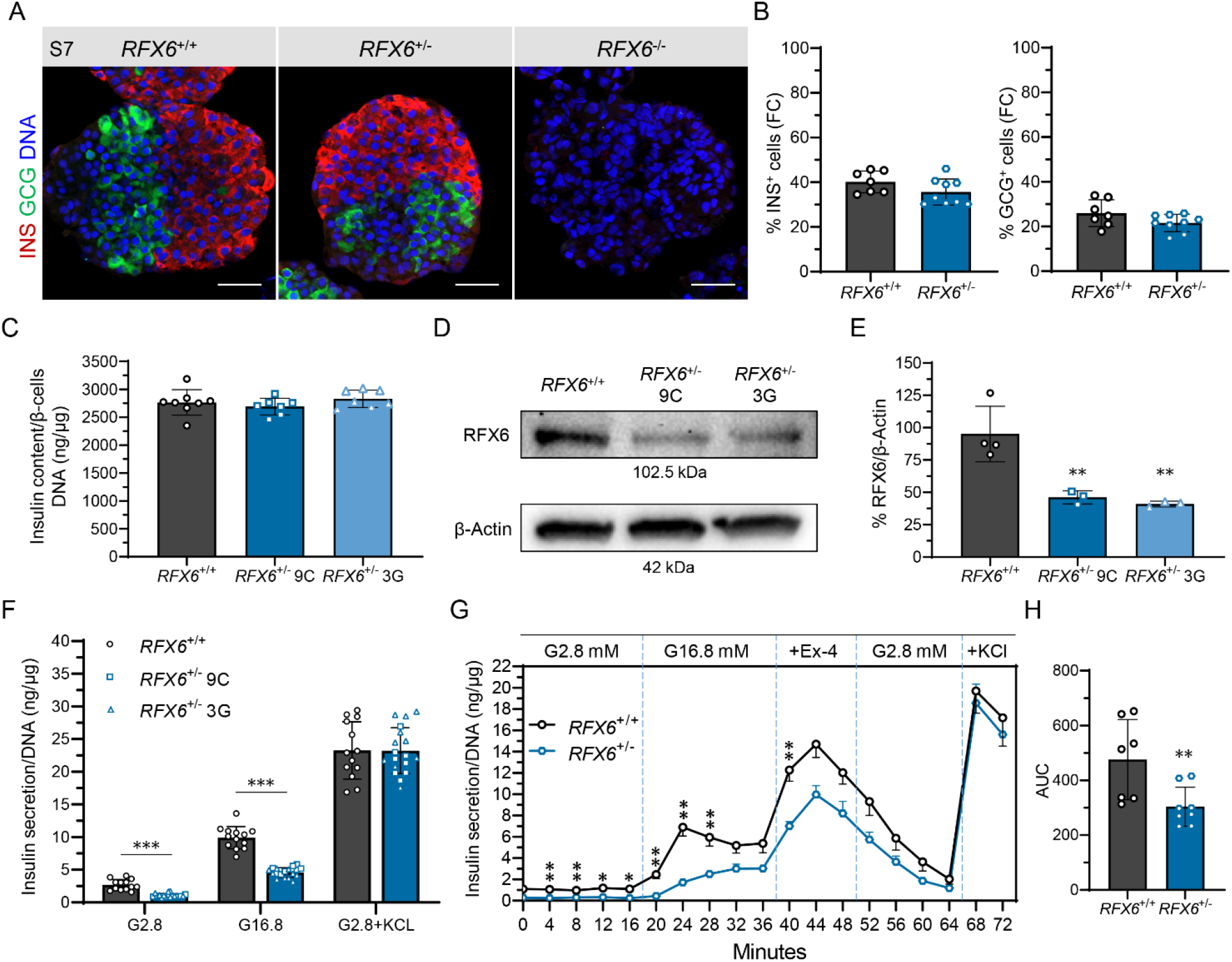
*RFX6* haploinsufficiency impairs insulin secretion of β-cells. **(A)** Immunohistochemistry showing GCG^+^ and INS^+^ cells for *RFX6*^+/+^ and *RFX6*^+/−^ SC-islets at S7w2, and for *RFX6*^−/−^ at S7d2, scale bars, 50 µm. **(B)** Percentage of INS^+^ and GCG^+^ cells for *RFX6*^+/+^ and *RFX6*^+/−^ SC-islets at S7w2 measured by flow cytometry (n=7–9). **(C)** Insulin content of *RFX6*^+/+^ and *RFX6*^+/−^ SC-islets at S7w3 normalized to the DNA content of the β-cells (n=7–8). **(D)** Immunoblot for RFX6 and β-Actin for *RFX6*^+/+^ and *RFX6*^+/−^ SC-islets at S7w2. **(E)** Percentage of RFX6 protein bands densitometry normalized to β-Actin bands (n=3–4). **(F)** Static insulin secretion at S7w3 at low 2.8 mM glucose (G2.8), followed by high 16.8 mM glucose (G16.8) and then depolarization at low glucose + 30 mM KCl, normalized to the DNA content of the SC-islets (n=13–18). **(G)** Dynamic insulin secretion responses to perifusion with 2.8 mM (G2.8) to 16.8 mM glucose (G16.8), 50 ng/ml exendin-4 (Ex-4) and 30 mM KCl. Normalized to the DNA content of the SC-islets (n=7–8). **(H)** Area under the curve (AUC) quantified from G (n=7–8). Statistical significance was measured using unpaired t-test in (B and H), using one-way ANOVA with Tukey’s test for multiple comparisons correction in (C and E) and using two-tailed unpaired multiple t-tests in (F and G). Error bars represent ±SD from the mean, *p<0.05, **p<0.01, ***p<0.001.

### Reduced [Ca^2+^]_i_ levels in basal and depolarized conditions in *RFX6*^+/−^ SC-islets

To mechanistically understand the reduced glucose sensitivity of *RFX6*^+/−^ SC-islets, we explored the stimulus-secretion coupling machinery of SC-islet β-cells by analyzing, cytoplasmic Ca^2+^ concentrations, ion channel conductance and exocytosis. The cytoplasmic Ca^2+^ concentration ([Ca^2+^]_i_) was recorded from individual cells in SC-islets loaded with Fura-2LR. In line with reduced basal insulin secretion, [Ca^2+^]_i_ at 3 mM glucose was significantly lower in *RFX6*^+/−^ than *RFX6*^+/+^ SC-islets (Fig. 5A, B). The [Ca^2+^]_i_ increase in response to 16.7 mM glucose was moderately reduced, but the difference did not reach statistical significance. Blockade of K_ATP_ channels with 1 mM tolbutamide increased [Ca^2+^]_i_ comparable to glucose stimulation. Depolarization with 30 mM K^+^ resulted in a more pronounced [Ca^2+^]_i_ increase, which was significantly reduced in *RFX6*^+/−^ compared to *RFX6*^+/+^ SC-islets (Fig. 5A, B). Patch-clamp recordings showed no difference in amplitude or voltage-dependence of total and nifedipine-sensitive Ca^2+^ currents or of Na^+^ currents (Supplementary Fig. 8C, D). Exocytosis was measured in single cells as changes in membrane capacitance. The pre-stimulatory capacitance was similar in control and *RFX6*^+/−^ cells, indicating no difference in cell size (Supplementary Fig. 8E). A train of depolarizations (14×200 ms) resulted in slightly larger capacitance increase in *RFX6*^+/−^ cells (Supplementary Fig. 8F, G), but the difference was not statistically significant. Exocytosis was also measured with total internal reflection fluorescence (TIRF) microscopy in cells expressing the granule marker NPY-tdmOrange2. Control and *RFX6*^+/−^ cells showed similar granule density at the plasma membrane and comparable degree of granule fusion events following depolarization with high K^+^ combined with diazoxide (Supplementary Fig. 8H, I). Together, these experiments indicate that *RFX6*^+/−^ SC islets have no general impairment of the exocytosis capacity and that the reduced glucose-stimulated insulin secretion is most likely explained by other factors, such as altered generation of metabolic coupling factors that amplify Ca^2+^-triggered exocytosis.

**Fig. 5.**
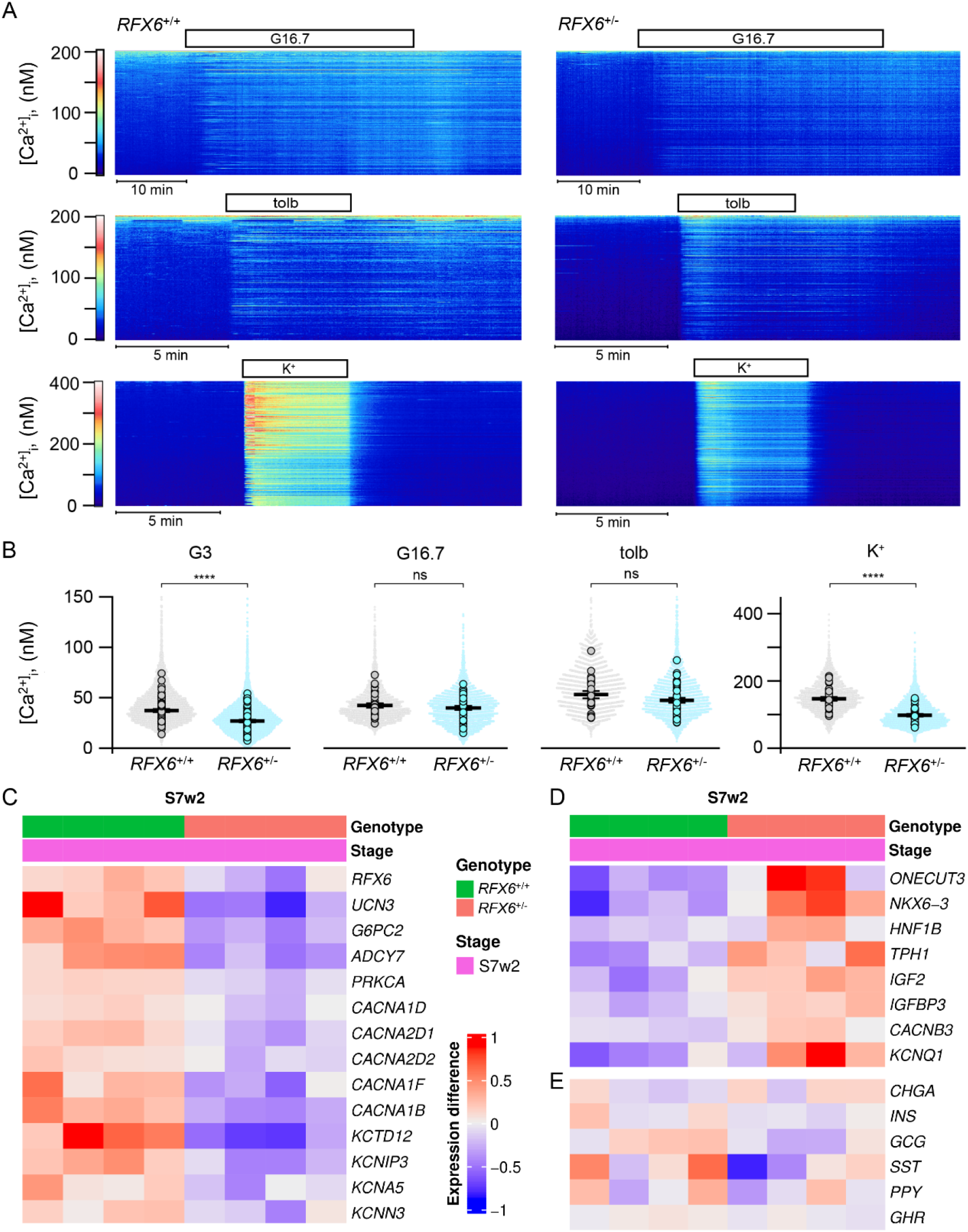
Reduced [Ca^2+^]_i_ levels in basal and depolarized conditions in *RFX6*^+/−^ SC-islets. **(A)** Heatmaps of [Ca^2+^]_i_ recorded with Fura-2 LR in cells within SC-islets under basal conditions (3 mM glucose) and after stimulation with 16.7 mM glucose, 1 mM tolbutamide and 30 mM K^+^. Each heatmap shows the responses of all cells in one islet. **(B)** Quantifications of [Ca^2+^]_i_ responses from experiments as in A. Time-averaged [Ca^2+^]_i_ for all individual cells (dots) and average values for all cells in each islet (markers). The bars represents means ±SEM for the averaged islet responses. n=number of islets (total cell number in parenthesis): G3 WT, n=65 (6933); G3 *RFX6*^+/−^, n=75 (8502); G16.7 WT, n=29 (3203); G16.7 *RFX6*^+/−^, n=42 (4853); tolb WT, n=23 (2720); tolb *RFX6*^+/−^, n=39 (4692); K^+^ WT n=32 (3639); K^+^ *RFX6*^+/−^, n=37 (4320). Statistical analyses with Student’s unpaired two-tailed t-test, ns; non-significant, ****p<0.0001. **(C)** Heatmap of downregulated genes in *RFX6*^+/−^ SC-islets from deep RNAseq analysis at stage 7 week 2 (n=4). **(D)** Similar heatmap of upregulated genes (n=4). **(E)** Heatmap of non-significantly different hormone gene expression (n=4). (C–E) Each gene is shown with a multiple testing corrected p value generated for the longitudinal differential expression of the gene during differentiation.

To further investigate the findings of lower insulin secretion and reduced cytoplasmic Ca^2+^ concentrations, we performed transcriptome analysis at S7 using bulk RNAseq (Supplementary Fig. 5). The gene expression levels of *RFX6* were downregulated in the heterozygous cells, in line with the observed lower protein expression and functional haploinsufficiency. Concomitantly, β-cell maturation markers such as, *G6PC2* (*28*), *UCN3* (*33*), *ADCY7* (*34*) and *PRKCA* (*35–37*), together with voltage-gated calcium and potassium channels, were reduced compared to the wild type (Fig. 5C). In contrast, pancreatic progenitor genes such as, *ONECUT3* (*38*), *NKX6.3* (*39*, *40*) and *HNF1B* (*41*) were upregulated (Fig. 5D), indicating an immature phenotype of the heterozygous SC-islets. Interestingly, the serotonin biosynthesis gene *TPH1* was increased, similar to what was found in Rfx6-deficient mice (*42*). Moreover, upregulation of the disallowed genes *IGF2* (*43*, *44*), *IGFBP3* (*45*, *46*), *CACNB3* (*47*, *48*) and *KCNQ1* (*49*, *50*) was detected (Fig. 5D). However, the gene expression of pancreatic hormones was unaltered (Fig. 5E). These findings illustrate the immature phenotype of *RFX6*^+/−^ SC-islets with reduced cytoplasmic Ca^2+^ levels and absolute insulin secretion.

### Heterozygous *RFX6* PTV β-cells sustain lower insulin secretion *in-vivo*

To investigate if the reduced insulin response persisted after further maturation of the SC-islets *in-vivo*, we implanted 500 S7 *RFX6*^+/+^ or *RFX6*^+/−^ SC-islets under the kidney capsule of non-diabetic NOD-SCID-Gamma (NSG) mice. Additionally, we implanted S6 *RFX6*^−/−^ SC-islets to evaluate the fate of the surviving cells *in-vivo*, since they failed to survive *in-vitro*. Blood glucose and human C-peptide levels were measured monthly for 3 months, followed by an intraperitoneal glucose tolerance test (IPGTT) (Fig. 6A). The normal range of blood glucose levels in random-fed NSG mice is between 6 and 8.5 mM, however, starting from the second month post-implantation, the blood glucose levels of the mice implanted with *RFX6*^+/+^ SC-islets dropped to the human level of ≈4.6 mM (Fig. 6B). In contrast, mice implanted with *RFX6*^+/−^ SC-islets required an additional month to display human blood glucose levels (Fig. 6B). *RFX6*^−/−^ cells failed to reduce the mouse blood glucose levels. While human C-peptide levels were undetectable in *RFX6*^−/−^ implanted mice, they increased steadily in the *RFX6*^+/+^ and *RFX6*^+/−^ implanted mice. However, throughout the 3 months, the C-peptide levels in *RFX6*^+/−^ mice were approximately half of their *RFX6*^+/+^ counterparts (Fig. 6C).

**Fig. 6.**
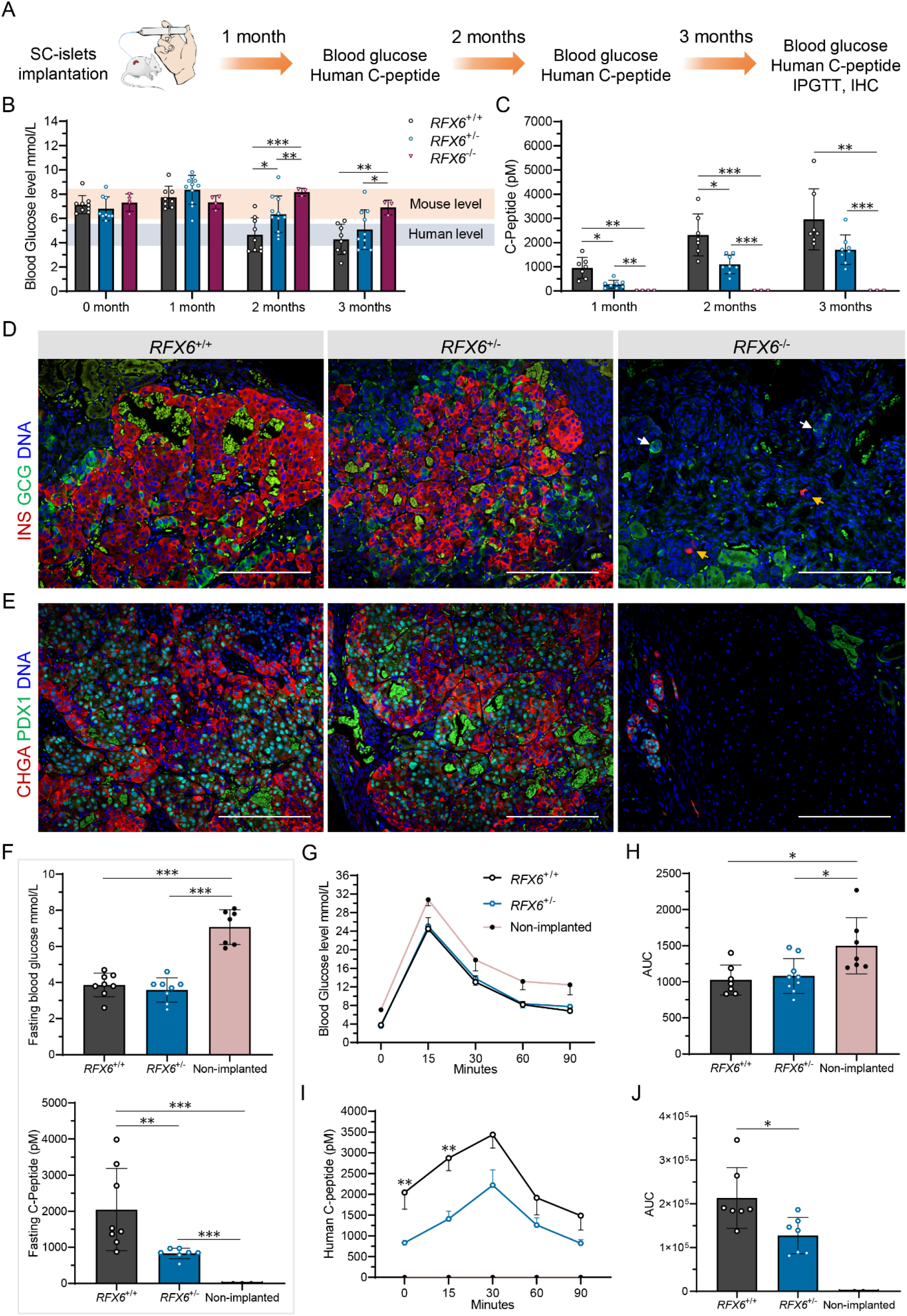
Heterozygous *RFX6*^+/−^ β-cells sustain lower insulin secretion *in-vivo*. **(A)** Schematic showing the data points and sampling post SC-islets implantation under the kidney capsule of NSG mice. **(B)** Random blood glucose levels measured monthly post implantation (n=3–11). **(C)** Random human C-peptide levels measured monthly post implantation (n=3–8). **(D)** Immunohistochemistry showing GCG^+^ and INS^+^ cells at month 3 post implantation. In *RFX6*^−/−^ panel, yellow arrows point to INS^+^ and white arrows point to GCG^+^, scale bars, 200 µm. **(E)** Immunohistochemistry showing CHGA^+^ and PDX1^+^ cells at month 3 post implantation, scale bars, 200 µm. **(F)** Upper panel shows fasting blood glucose levels measured at month 3 post implantation (n=7– 8). Bottom panel shows fasting human C-peptide levels measured at month 3 post implantation (n=3–8). **(G)** Blood glucose levels during an IPGTT of fasted mice at month 3 post implantation (n=7–9). **(H)** Area under the curve (AUC) quantified from F (n=7–9). **(I)** Human C-peptide during an IPGTT of fasted mice at month 3 post implantation (n=2–7). **(J)** Area under the curve (AUC) quantified from H (n=2–7). Statistical significance was measured using two-way ANOVA with Tukey’s test for multiple comparisons correction in (B and C), using one-way ANOVA with Tukey’s test for multiple comparisons correction in (F–I) and using two-tailed unpaired t-test between *RFX6*^+/+^ and *RFX6*^+/−^ in (J). Error bars represent ±SD from the mean except for (G and I) showing ±SEM, *p<0.05, **p<0.01, ***p<0.001.

Immunohistochemical analysis after 3 months showed abundant INS^+^ and GCG^+^ cells in *RFX6*^+/+^ and *RFX6*^+/−^ grafts, while these cell types were rarely found in *RFX6*^−/−^ implants (Fig. 6D). Similarly, majority of the implanted cells in *RFX6*^+/+^ and *RFX6*^+/−^ were CHGA^+^ with an abundance of PDX1^+^ cells, representing INS^+^ cells. In contrast, the majority of cells were negative for both markers in *RFX6*^−/−^ implants and the morphology of cells was distinctly different (Fig. 6E). To evaluate the origin of these cells, we performed immunohistochemical analysis using a human-specific mitochondrial antibody (hMITO), together with SOX9 immunostaining that was sustained in *RFX6*^−/−^ *in-vitro*. Indeed, INS^+^ cells from the *RFX6*^+/+^ implants were hMITO^+^, while the surrounding cells from the mouse tissue were negative (Supplementary Fig. 9A). The morphologically different cells in the *RFX6*^−/−^ implants were hMITO^+^ and were positive for nuclear SOX9, compared to scarce cytoplasmic SOX9^+^ cells in *RFX6*^+/+^ implants. Both *RFX6*^+/+^ and *RFX6*^−/−^ cells were negative for NEUROG3 expression at 3-months post-implantation (Supplementary Fig. 9B). Since PP cells were found in *Rfx6*^−/−^ mice, we stained for PP and did not detect any PP^+^ cells in the *RFX6*^−/−^ grafts (Supplementary Fig. 9C). This suggests that RFX6 is indispensable for the development of all pancreatic endocrine cells including PP cells in humans.

At 3 months of implantation *RFX6*^+/+^ and *RFX6*^+/−^ implanted mice had a blood glucose level of ≈3.9 mM, similar to fasting human level, while non-implanted mice presented the normal mouse blood glucose levels around ≈7 mM (Fig. 6F). However, human C-peptide levels in fasting mice with *RFX6*^+/−^ SC-islets were ≈60% lower (Fig. 6F). In intraperitoneal glucose tolerance test (IPGTT), both *RFX6*^+/+^ and *RFX6*^+/−^ SC-islets were able to control the abrupt increase in blood glucose levels similarly, and more efficiently compared to the non-implanted mice, which had higher blood glucose levels in each of the measured time points (Fig. 6G) and in the measured AUC (Fig. 6H). Although the secreted human C-peptide levels of both genotypes had a similar pattern and peaked at 30 min post-injection, the absolute C-peptide secretion was lower in the *RFX6*^+/−^ implanted mice (Fig. 6I) and the total secretion AUC was significantly reduced (Fig. 6J). These findings indicate that RFX6 haploinsufficiency due to the lower gene dose in *RFX6*^+/−^ SC-islets leads to sustained reduction in insulin secretion.

## Discussion

A hierarchy of gene regulatory networks control the development and function of β-cells. Single pathogenic variants in more than twenty transcription factors of this network can lead to diabetes. Importantly, species differences in the developmental and functional processes of β-cells impede their precise study with mouse models. Here, we sought to overcome this limitation using genetically-engineered human stem-cell-derived pancreatic endocrine cells (SC-islets) to investigate the developmental and functional defects of β-cells upon *RFX6* perturbation.

Our findings demonstrate that RFX6 p.His293LeufsTer7 is a loss-of-function PTV with a transcript subjected to nonsense-mediated mRNA decay. We confirmed that the lower *RFX6* gene dosage in heterozygous *RFX6* SC-islets led to *RFX6* haploinsufficiency, which did not compromise the differentiation capacity of the cells in terms of β-cell number or insulin content, but impaired β-cell functionality. There was no association between *Rfx6* haploinsufficiency and diabetes in mice, indicating that this phenotype cannot be faithfully modelled in rodents (*6*). Similar discrepancies between diabetes development in humans and mice have been reported with heterozygous mutations in *HNF1A*, *HNF1B* and *PAX4*. While pathogenic heterozygous mutations in all of these genes cause MODY in humans, heterozygous null mice lack any diabetes phenotype (*51–53*). However, our findings are in line with the β-cell-specific *Rfx6* knockout in adult mice (*12*). Reduced RFX6 expression in *RFX6^+/−^* SC-islets led to decreased levels of the β-cell marker *UCN3* and VDCCs expression resulting in lower cytoplasmic Ca^2+^ concentrations, reducing absolute insulin secretion in basal and high glucose levels. In contrast, *RFX6^+/−^* SC-islets showed upregulation of disallowed genes that are reported to negatively impact insulin secretion, such as *IGF2* (*43*, *44*), *IGFBP3* (*45*, *46*), *CACNB3* (*47*, *48*) and *KCNQ1* (*49*, *50*).

Insulin content of the heterozygous cells was not affected, suggesting that the difference in insulin secretion more likely reflects impaired stimulus-secretion coupling or granule trafficking. However, exocytosis measured either as changes in capacitance or by TIRF imaging of granules at the plasma membrane was not reduced in response to depolarizing stimuli, which is in line with intact K^+^-induced insulin secretion in static and dynamic perifusion experiments. Despite slightly reduced gene expression levels of some voltage-gated Ca^2+^ channel components, there was no difference in Ca^2+^ currents, unlike the reduced Ca^2+^-channel activity that was reported in *RFX6* knockdown in EndoC-βH2 (*13*). The reduced basal [Ca^2+^]_i_ and the lower [Ca^2+^]_i_ after membrane depolarization may instead reflect differences in Ca^2+^ transport or buffering. The lower basal [Ca^2+^]_i_ is in itself consistent with reduced insulin secretion at low glucose. In addition, substimulatory cytosolic [Ca^2+^]_i_ also promotes granule priming (*54*), and may thereby indirectly amplify secretion at elevated glucose. However, the reduced insulin secretion despite insignificantly altered [Ca^2+^]_i_ at high glucose more likely points towards a role for *RFX6* in controlling the metabolic amplifying pathway. We suggest insulin secretagogues or GLP-1 receptor agonists as a treatment to enhance the secretory capacity of these β-cells.

The heterozygous *RFX6* SC-islets sustained lower insulin secretion *in-vivo*, resulting in delayed lowering of blood glucose post-implantation. All of our findings on the heterozygous cell model are consistent with the increased risk of variant carriers to develop diabetes, as shown for both gestational diabetes and T2D, and previously for MODY with reduced penetrance (*15*). The high carrier frequency and the subtle albeit significant clinical differences in presentation in the FinnGen cohort do not suggest marked insulin-deficiency in heterozygote carriers nor capability for causing monogenic diabetes. The clinical phenotype of the variant carriers needs more exploring, but two small studies have shown higher fasting (*15*) or 2-hour glucose (*30*) during an oral glucose tolerance test in carriers compared to non-carriers, as well as lower fasting and stimulated GIP level (*15*). In agreement with an effect on insulin secretion, the BMI and age at diagnosis of diabetes was younger in the carriers.

The complete loss of RFX*6* led to reduced expression of *PDX1*, *NKX6.1*, *FEV*, *ISL1, NEUROD1* and *PAX6* with gradual loss of pancreatic progenitor identity. This is in line with some of the reported targets of ChIP-seq analysis of whole pancreatic tissue from adult mice such as *Pdx1*, *Neurod1* and *Nkx6-1* (*55*). Although, the pancreatic markers *NEUROG3*, *NKX2.2* and *PAX4* were reduced at the posterior foregut stage, they recovered at the endocrine precursor stage, suggesting a negative feedback loop between RFX6 and NEUROG3 at this stage. This phenotype is similar to what was shown in *Rfx6* knockout mice (*6*, *12*), with a difference of reduced *NKX6.1* expression in our model. While gene expression levels of *PPY* were increased in *RFX6*^−/−^ endocrine precursors, PP cells did not form *in-vitro* or after 3 months of implantation *in-vivo*, recapitulating the loss of pancreatic polypeptide in the Mitchell-Riley syndrome patient, which is distinct from the presence of PP cells in the mouse model (*6*). We showed sustained NEUROG3 and SOX9 expression after the pancreatic endocrine stage in *RFX6*^−/−^ cells, indicating an endocrine specification barricade, which was followed by increased cell death when the cells were directed to further differentiate *in-vitro*. The reduced generation of pancreatic progenitor pool and the increased apoptosis phenotype could explain the pancreatic hypoplasia seen in patients. Implantation of the homozygous cells, confirmed sustained SOX9 expression in the majority of grafted cells, while CHGA expressing cells were extremely rare. This suggests that RFX6 functions to limit SOX9 expression to allow full endocrine differentiation. We did not detect human C-peptide in mice implanted with *RFX6*^−/−^ cells, however, the presence of small numbers of INS^+^ cells and GCG^+^ cells may explain the very low C-peptide levels detected in the patient, but not the normal glucagon levels. The normal glucagon levels in the patient might be explained by extrapancreatic secretion of the hormone (*56*).

In summary, we highlight the critical role of RFX6 in augmenting and maintaining the pancreatic progenitor pool, with an endocrine roadblock upon its loss, sustaining NEUROG3 and SOX9 expression and increasing cell death. We demonstrate that *RFX6* haploinsufficiency does not affect β-cell number or insulin content, but does impair functionality, predisposing carriers to diabetes. Our allelic series isogenic SC-islet models represent a powerful tool to elucidate specific etiologies of diabetes in humans, enabling the sensitive detection of aberrations in both β-cell development and function.

## Materials and Methods

### Lifetime risk analysis of T2D and gestational diabetes in the FinnGen dataset

We analyzed the risk of gestational or T2D, the age at diagnosis of diabetes and body-mass index (BMI) in the *RFX6* variant carriers compared to non-carriers in the FinnGen data freeze R11 at individual level, using genome-wide genotype data and longitudinal healthcare registry data (*23*, *57*). FinnGen (https://www.finngen.fi/en) is a public-private research project, combining genome and digital healthcare data since 2017. In FinnGen, diseases are defined as endpoints by FinnGen clinical expert teams according to ICD codes of version 8 (1969–1986), 9 (1987–1995), and 10 (1996–2019). The FinnGen partners can be found at https://www.finngen.fi/en/partners. FinnGen R11 comprises 440,734 Finnish European individuals with informed consent for biobank research based on the Finnish Biobank Act, or regarding earlier cohorts, with approval from Fimea, the National Supervisory Authority for Welfare and Health. The FinnGen study protocol (number HUS/990/2017) is approved by the Coordinating Ethics Committee of the Hospital District of Helsinki and Uusimaa (HUS).

### Generating Mitchell-Riley syndrome patient-derived iPSCs and stem cell culturing

Dermal fibroblasts obtained from a skin biopsy from the patient at 6 months of age were reprogrammed using Sendai viruses (SeVdp), containing OCT4, SOX2, KLF4, and C-MYC reprogramming factors. The formed HEL118 iPSC lines were verified using Sanger sequencing to confirm the *RFX6* gene variant, were tested negative for mycoplasma contamination and did not contain transgene vectors. Additionally, the iPSCs line HEL46.11 (derived from human neonatal foreskin fibroblast) and the Human embryonic stem cell line H1 (WA01, WiCell), which are wild-type *RFX6* were used in this study. All the generated stem cell-lines were cultured on Matrigel-coated (Corning, #354277) plates with E8 medium (Thermo Fisher, A1517001) and passaged using 0.5 mM EDTA (Life Technologies, 15575–038) in PBS as a dissociation agent. The patient iPSCs were derived after informed consent and according to the approval of the coordinating ethics committee of the Helsinki and Uusimaa Hospital District (no. 423/13/03/00/08). The patient clinical data was used after informed consent.

### Genome editing of iPSCs and H1 hESCs

For correcting the *RFX6* PTV in the patient’s HEL118.3 iPSCs, we used a plasmid-based CRISPR-Cas9 system with a guide and a correction template that were designed with Benchling (Biology Software, 2017) (gRNA1 correction: ATAACAGGATTTTCGAGCAG). The correction template introduced a TseI restriction site by silent mutations to facilitate the screening of the edited clones. Two million iPSCs were electroporated with 6 µg of CAG-Cas9-T2A-EGFP-ires-Puro (deposited in Addgene, plasmid no. 78311, together with detailed protocols for its use), 500 ng of the gRNA cassette and 4 µg of the 200 bp dsDNA correction template, using Neon Transfection system (Thermo Fisher Scientific; 1100 V; 20 ms; two pulses). Single cells were sorted, expanded, and screened using TseI restriction enzyme. The editing resulted in heterozygous clones only, the heterozygous clone 10B was electroporated using gRNA2 correction guide (AGCAGGGGAAGGAGATGGTC) to obtain the homozygous clone 2D. Positive clones were further validated by Sanger sequencing.

For introducing the *RFX6* PTV in H1 hESCs, we used a more efficient Ribonucleoprotein (RNP) CRISPR-Cpf1 system (Integrated DNA Technologies), with a guide and a variant template that were designed with Benchling (Biology Software, 2017) (gRNA mutation: TTACACTTTTGGCAAGGAATG). Two million cells were electroporated with 10 µg Alt-R™ A.s. Cas12a (Cpf1) Ultra combined with the gRNA and 4 µg of the 100 b ssODN correction template (Integrated DNA Technologies), using Neon Transfection system (Thermo Fisher Scientific; 1100 V; 20 ms; two pulses). Single cells were sorted, expanded, and screened using TseI restriction enzyme. The editing resulted in heterozygous clones only, the heterozygous clone 3G was electroporated with the same gRNA and a mutation template carrying TseI and NheI restriction sites to facilitate the screening of homozygous clones. Positive clones were further validated by Sanger sequencing. The top three CRISPR off-target hits were sequenced and did not have any indels. Karyotype analyses based on chromosomal G-banding were performed for all the generated hiPSCs and hESCs clones at Ambar Lab, Barcelona, Spain. Primer sequences used for genome editing are described in Supplementary Table 1.

### *In-vitro* pancreatic endocrine differentiation culture

HEL118.3 iPSCs were differentiated towards the pancreatic lineage to generate pancreatic endocrine cells using the previously published detailed protocol with minor modifications. Briefly, two million cells/well were seeded on 6-well Matrigel coated plates in E8 medium supplemented with 10 µM Rho-Associated kinase inhibitor (ROCKi, Y-27632, Selleckchem S1049). The differentiation was started following 24h of seeding and proceeded through a 6-stage differentiation protocol in monolayer. The hESCs H1 cell-lines were differentiated using the extended 7-stage protocol (stages 1–4 in adherent culture, stage 5 in AggreWell (Stemcell Technologies, #34421), and stages 6 and 7 in suspension culture).

The detailed differentiation protocol and the stage specific complete media formulations are described in Supplementary Table 2 and 3. The details of the reagents used in the differentiation protocol are described in Supplementary Table 4.

### Western blot

For protein extraction, cells were washed with ice-cold PBS and lysed with Cell lysis buffer (Cell Signaling Technologies #9803) for 10 min on ice. The cells were sonicated for 3 x 5 s on ice, centrifuged (10000 rcf for 10 min at 4°C) and the supernatant was stored at −80°C. The samples were run on Any kD™ Mini-PROTEAN^®^ TGX™ gel (Bio-Rad Laboratories) and then dry transferred onto a nitrocellulose membrane using the iBlot system (Invitrogen) as per manufacturer’s instructions. The membrane was then probed with the primary antibody overnight at 4°C, washed twice with Tris-buffered saline (Medicago #09-7500) containing 0.1% Tween for 2 x 10 min, and incubated with the corresponding secondary antibody for 30 min at room temperature. Chemiluminescence detection was performed with Amersham ECL (RPN2235; Cytiva) and Bio-RAD Chemidoc XRS1 imaging system; Image Lab software v6.0.0. Densitometry quantification analysis was done using Fiji v1.53. The details of antibodies and their dilutions used for WB are described in Supplementary Table 5.

### Flow cytometry

For quantifying the percentage of definitive endoderm positive cells after stage 1, flow cytometry for CXCR4 was performed on one million cells of dissociated cells using TrypLE (Thermo Fisher Scientific) for 3 min at 37°C. Cells were resuspended in cold 5% FBS-containing PBS and the conjugated antibodies were added and incubated for 30 min at RT. For intracellular antigen cytometry of stages 4, 5, 6 and 7, cells were dissociated with TrypLE for 8 min at 37°C and resuspended in cold 5% FBS-containing PBS. Cells were fixed and permeabilized using Cytofix/Cytoperm (554714, BD Biosciences) as per manufacturer’s instructions. Primary or conjugated antibodies were incubated with the cells overnight at 4°C in Perm/Wash buffer (554714, BD Biosciences) containing 4% FBS and then secondary antibodies for 45 min at RT. The cells were then analyzed using FACSCalibur cytometer (BD Biosciences) with BD Cellquest Pro v4.0.2 (BD Biosciences) and FlowJo (Tree Star Inc.) softwares. The details of antibodies and their dilutions for flow cytometry are described in Supplementary Table 5.

### Immunocytochemistry and immunohistochemistry

For whole mount or adherent cultures staining, cells were fixed in 4% PFA for 15 min at RT, permeabilized with 0.5% Triton X-100 in PBS for 15 min at RT, then blocked with Ultra-V (Thermo Fisher Scientific) for 10 min and incubated with primary antibodies overnight at 4°C and secondary antibodies for 1h at RT diluted in 0.1% Tween in PBS. For paraffin embedding, aggregates were fixed with 4% PFA at 4°C for 24h, aggregates were embedded in 2% low-melting agarose (Fisher Bioreagents) PBS and transferred to paraffin blocks. Implanted grafts were retrieved after 3 months, dissected, and fixed with 4% PFA at RT for 48h, paraffin embedded, and cut into 5 µm sections using Leica microtome. For immunohistochemistry, slides were deparaffinized and antigens retrieved by boiling slides in 0.1 M citrate buffer (pH 6) using Decloaking chamber (Biocare Medical) at 95°C for 12 min. For TUNEL assay, In Situ Cell Death Detection Kit, Fluorescein (#11684795910; Roche) was used according to the manufacturer’s instruction. Invitrogen™ EVOS™ FL Digital Inverted Fluorescence Microscope or Zeiss Axio Observer Z1 with Apotome were used to image the cells and further processed with ZEN-2 software. All stained samples were equally treated and imaged with the same microscope parameters. Image quantification was performed using CellProfiler software v4.2.1 and Fiji software v1.53f. The details of antibodies and their dilutions used in the study for immunofluorescence are described in Supplementary Table 5.

### RNA extraction and quantitative RT-PCR

Total RNA was extracted using NucleoSpin Plus RNA kit (Macherey-Nagel). A total of 1.5 µg RNA was reversely transcribed using 0.5 μl Moloney murine leukemia virus reverse transcriptase (M1701, Promega), 4 µl RT buffer 5x (Promega), 2.5 μl dNTPs 2.5 mM, 1 μl Oligo-dT 500 μg/ml (Promega), 0.2 μl Random hexamers 500 μg/ml (Promega) and 0.5 μl Riboblock RNase inhibitor 40 u/μl (Fermentas) for 90 min at 37°C. The generated cDNA was amplified using 5x HOT FIREPol EvaGreen qPCR Mix Plus no ROX (Solisbiodyne) in a 20 µl reaction. The reactions were pipetted using QIAgility (Qiagen) robot into 100 well disc run in Rotor-Gene Q. Relative quantification of gene expression was analysed using ΔΔCt method, with cyclophilin G (PPIG) as a reference gene. Reverse transcription without template was used as negative control and exogenous positive control was used as a calibrator. The qRT-PCR primers sequences are described in Supplementary Table 6.

### NMD cycloheximide inhibition assay and NGS sequencing

At stage 3 of the differentiation protocol, heterozygous corrected HEL118.3 *RFX6*^+/−^ cells were treated with different concentrations of cycloheximide (Sigma-Aldrich, 01810) (25, 50 and 100 µg/ml) for durations of 3 and 5 h. Total RNA was collected and reversely transcribed as described in the previous section, but without using random hexamers. Primers with illumina sequences attached were designed and used to amplify 120 bp in the variant region of the *RFX6* cDNA, and 10 µl (5 ng/µl) were sequenced using NGS sequencing HiSeq PE100. The percentages of variant and corrected filtered reads from each of the alleles were calculated and plotted for the non-treated control and the cycloheximide treated samples. Primer sequences used for NGS cDNA amplification are described in Supplementary Table 1.

### Ultra-deep bulk RNAseq analysis

We performed ultra-deep RNAseq analysis for H1 *RFX6*^+/+^ and *RFX6*^−/−^ at stage 3 (PF) and stage 5 (EP), and for *RFX6*^+/+^ and *RFX6*^+/−^ at stage 7 (SC-islets). Following mRNA library preparation using NEBNext Ultra II Directional RNA kit, sequencing was performed using NovaSeq SP 2×100 bp v1.5 chemistry. Preparation of RNA library and transcriptome sequencing was performed by the Sequencing laboratory of Institute for Molecular Medicine Finland FIMM Technology Centre, University of Helsinki.

The raw data was filtered with cutadapt (*58*) to remove adapter sequences (with a minimum overlap of 5bp), ambiguous (N) and low-quality bases (Phred score < 25). We also excluded read pairs that were too short (<25bp) after trimming. The filtered read pair were mapped to the human reference genome (GRCh38) with STAR aligner (*59*). Gene expression was counted from read pairs mapping to exons using featureCounts in Rsubreads (*60*), using GENCODE (GRCh38.p13) genome annotations (*61*). Duplicates, chimeric and multimapping reads were excluded, as well as reads with low mapping scores (MAPQ < 10). We obtained on average 31M uniquely mapped read pairs per sample (between 27-38M).

We removed genes with very low or no expression, such as non-polyadenylated genes, from the analysis (<50 reads across all samples). The read count data were analyzed with DESeq2 (*62*), comparing the expression differences between knockout and control samples in the different developmental stages. For stage 3 and 5 we compared the homozygous *RFX6*^−/−^ variant (clone 1H) to the control *RFX6*^+/+^ (clone 10F) samples. Since *RFX6*^−/−^ cells failed to differentiate into endocrine lineage, we compared the heterozygous *RFX6*^+/−^ (clone 3G) to the control *RFX6*^+/+^ (clone 10F) samples at stage 7 (week 2). Each genotype per stage had 4 biological replicates.

Genes were considered differentially expressed if the False Discovery Rate (FDR) adjusted *p* values were <0.01. PCA was calculated with prcomp using normalized, log1p-transformed read counts. The differentially expressed genes (FDR<0.01) were analyzed for enrichment separately for the up- and down-regulated genes using clusterProfiler (*63*) against Reactome (*64*), KEGG (*65*), Disease ontology (*66*) and Gene ontology (*67*) databases. For heatmaps we used log1p-transformed normalized read counts, centered around gene mean using ComplexHeatmap (*68*). For the Venn diagram we used ggvenn. The rest of the RNA-seq figures were made with ggplot2 (*69*). The results from the gene expression analysis together with the raw sequences were deposited to GEO, under accession GSE234289.

## STAR Methods

Fastq filtering: cutadapt (version 2.6)

Mapping / alignment software: STAR (version 2.7.6a)

RNA-seq analysis software: R (version 4.1) using Bioconductor (version 3.14)

Read counting: Rsubread (version 2.8.2)

Venn diagram: ggvenn (version 0.1.8) https://github.com/yanlinlin82/ggvenn

Heatmap figures: ComplexHeatmap (version 2.10.0)

Volcano plot, Boxplot, and PCA figures: ggplot2 (version 3.3.5)

Genome annotations: GENCODE GRCh38.p13

RNA-seq adapter sequences:

5’-AGATCGGAAGAGCACACGTCTGAACTCCAGTCA-3’ (R1),

5’-AGATCGGAAGAGCGTCGTGTAGGGAAAGAGTGT-3’ (R2)

### Static and dynamic glucose-stimulated insulin secretion

For the static glucose-stimulated insulin secretion assay, fifty aggregates were preincubated in 2.8 mmol/L glucose Krebs buffer in a 12-well plate placed on a 95-rpm rotating platform for 90 min at 37°C. Aggregates were then washed with Krebs buffer and sequentially incubated in Krebs buffer containing 2.8 mmol/l glucose, 16.6 mmol/l glucose, and 2.8 mmol/l glucose plus 30 mmol/l KCl, for periods of 30 min each. Samples of 200 µl were collected from each treatment and stored at −80°C for insulin ELISA measurements. Dynamic tests of insulin secretion were carried out using a perifusion apparatus (Brandel Suprafusion SF-06, MD, USA) with a flow rate of 0.25 ml/min and sampling every 4 minutes. Samples from each fraction collected were analyzed using insulin ELISA (Mercodia, Sweden). Following static and dynamic tests of insulin secretion, the SC-islets were collected, and the total insulin and DNA contents were analyzed. Stimulated insulin secretion results are normalized using DNA content of the β-cell fraction, which was calculated from flow cytometry INS^+^ cell percentage.

### Electrophysiology

SC-islets were dispersed into single cells in cell dissociation buffer (Thermo Fisher Scientific) supplemented with trypsin (0.005%, Life Technologies), washed and plated in serum-containing medium on 22-mm polylysine-coated coverslips, allowed to settle overnight, and then transduced with adenovirus coding for enhanced green fluorescent protein under control of the RIP2 promoter to identify beta cells. Patch-clamp recordings were performed using an EPC-9 patch amplifier with PatchMaster v.2×90 software (HEKA Electronics). Electrodes (resistance 2–4 MΩ) were pulled from borosilicate glass capillaries, coated with Sylgard and fire-polished. Cells were superfused with an extracellular solution containing 138 mM NaCl, 5.6 mM KCl, 1.2 mM MgCl_2_, 2.6 mM CaCl_2_, 10 mM glucose, and 5 mM HEPES, pH 7.4 adjusted with NaOH at a rate of 0.4 ml min^−1^ at 32°C. Voltage-dependent currents and exocytosis were measured in whole-cell voltage-clamp mode with an intracellular solution containing 125 mM Cs-glutamate, 10 mM CsCl, 10 mM NaCl, 1 mM MgCl_2_, 0.05 mM EGTA, 3 mM Mg-ATP, 0.1 mM cAMP and 5 mM HEPES, pH 7.2 adjusted using CsOH. For current-voltage (IV) relationships, the membrane was depolarized from −70 mV to +80 mV (in 10 mV steps) lasting 50 ms each. Currents were compensated for capacitive transients and linear leak using a P/4 protocol. Na^+^ and Ca^2+^ current components were separated by quantifying the initial peak current (0–5 ms; Na^+^) and average sustained current (5–45 ms; Ca^2+^). Exocytosis was quantified using the lockin module of Patchmaster (30 mV peak-to-peak; 1 kHz); with a train of 14 × 200 ms depolarizations to 0 mV at 1.4 Hz.

### Exocytosis imaging

To visualize granule exocytosis, cells treated as described for electrophysiology were additionally infected with adNPY-tdOrange2 (a well-established marker for secretory granules) and imaged after 30–36 h using a custom-built lens-type TIRF microscope based on an AxioObserver Z1 with a ×100/1.45 objective (Zeiss). Excitation was from two DPSS lasers at 491 and 561 nm (Cobolt) passed through a cleanup filter (catalog no. zet405/488/561/×640x; Chroma) and controlled with an acousto-optical tunable filter (AA-Opto). Excitation and emission light were separated using a beamsplitter (catalog no. ZT405/488/561/640rpc; Chroma). The emission light was separated chromatically onto separate areas of an EMCCD camera (Roper QuantEM 512SC) using an image splitter (Optical Insights) with a cutoff at 565 nm (catalog no. 565dcxr; Chroma) and emission filters (catalog nos. ET525/50 m and 600/50 m, Chroma). Scaling was 160 nm per pixel. Cells were imaged in a standard solution containing 138 mM NaCl, 5.6 mM KCl, 1.2 mM MgCl_2_, 2.6 mM CaCl_2_, 10 mM glucose, 5 mM HEPES (pH 7.4 with NaOH), 200 μM diazoxide (to block spontaneous depolarization; Sigma-Aldrich), and 10 nM exendin-4 (a GLP-1 receptor agonist; Anaspec). Exocytosis was evoked by rapidly depolarizing cells with elevated K^+^ (75 mM KCl equimolarly replacing NaCl in the standard solution, by computer-controlled local pressure ejection).

### [Ca^2+^]_i_ imaging

SC-islets were loaded with the fluorescent indicator Fura-2 LR (ion Biosciences) by 1 h incubation with 1 μM of its acetoxymethyl ester at 37°C in experimental buffer containing 138 mM NaCl, 4.8 mM KCl, 1.2 mM MgCl_2_, 2.56 mM CaCl_2_, 3 mM glucose, 25 mM HEPES (pH set to 7.40 with NaOH) and 0.5 mg ml^−1^ BSA. After rinsing in indicator-free buffer, the islets were attached to poly-l-lysine-coated coverslips in a 50-μl chamber on the stage of an Eclipse TE2000U microscope (Nikon) and superfused with buffer at a rate of 160 μl min^−1^. The chamber holder and ×40, 1.3-NA objective were maintained at 37°C by custom-built thermostats. An LED light engine (LedHUB, Omicron Laserage Laserprodukte) equipped with 340 and 385 nm diodes and 340/26 nm (center wavelength/half-bandwidth) and 386/23 nm interference filters (Semrock, IDEX Health & Science, LLC) provided excitation light that was led to the microscope via a liquid light guide. Emission was measured at 510/40 nm using a 400 nm dichroic beamsplitter and an Evolve 512 EMCCD camera (Photometrics). Image pairs at 340/386 nm were acquired every 2 s with the MetaFluor v.7.7 software (Molecular Devices). [Ca^2+^]i was calculated from the background-corrected Fura-2 LR 340/380 nm fluorescence excitation ratio from manually defined cell-sized regions of interest. The data are presented as heatmaps from individual islets and cells and as scatter plots of the absolute [Ca^2+^]_i_ values under different conditions. Calibration of Fura-2LR signal was performed to linearize the ratio signal using solutions without (0 Ca^2+^, 2 mM EGTA) and with saturating Ca^2+^ (10 mM) and 100 µM Fura-2LR salt. The fluorescence ratio was converted to [Ca^2+^]_i_ as described by Grynkiewicz *et al.* (*70*).

### *In-vivo* animal implantation studies

Animal care and experiments were approved by National Animal Experiment Board in Finland (ESAVI/14852/2018). NOD-SCID-Gamma (NSG, 005557, Jackson Laboratory) mice were obtained from SCANBUR and housed at Biomedicum Helsinki animal facility, on a 12h light/dark cycle and fed standard chow ad libitum, 2016 Teklad global 16% protein rodent diets (ENVIGO). The temperature was kept at 23°C with 24 relative humidity (RH). Implantations were performed on 4- to 8-month-old mice. Briefly, stage-6 *RFX6*^−/−^, and stage-7 *RFX6*^+/+^ and *RFX6*^+/−^ SC-islets equivalent to 1.5 million cells (500 clusters) were loaded in PE-50 tubing and implanted under the kidney capsule. Mouse serum samples were collected monthly from the saphenous vein and stored at −80°C for human C-peptide analysis. Blood glucose levels were measured using Contour^®^XT and Contour^®^next strips. Human-specific C-peptide was measured from plasma samples with the Ultrasensitive C-peptide ELISA kit (Mercodia, Uppsala, Sweden). To test the functionality of the SC-islet grafts, mice were subjected to an intraperitoneal glucose tolerance test (IPGTT). Mice were fasted 8 h before the test. The mice were weighed, and blood glucose was measured before the test. Glucose (3 g/kg) was injected intraperitoneally, and blood samples (30 μl) were taken from the saphenous vein after 15, 30, 60 and 90 min to measure blood glucose and circulating human C-peptide levels by ELISA.

### Quantification and statistical analysis

Data are collected from at least four independent differentiation experiments. Blinding was applied for immunohistochemical quantification. Morphological data represents population-wide observation from independent differentiation experiments. All the representative images in the figures were reproducible in all the experiments performed and are followed by a quantification panel for all the experiments. Bar graphs are represented showing all the individual data points. Statistical methods used are described in each figure legend and individual method section. The results are presented as the mean ±S.D unless otherwise stated. P-values<0.05 were considered statistically significant (*p<0.05; **p< 0.01; ***p<0.001).

### Data availability

Ultra-deep bulk RNAseq data for pancreatic differentiation stages −3, −5 and −7 of H1 *RFX6* genotypes are deposited in the Gene Expression Omnibus database with accession code (GSE234289). Original western blot images are deposited at Mendeley (DOI: 10.17632/g75drr3mgw.1). Any additional information required to reanalyse the data reported in this paper is available from the lead contacts on request, Hazem Ibrahim and Timo Otonkoski.

## Acknowledgements and funding

We thank Heli Grym, Anni Laitinen and Ras Trokovic for their expert technical support. T.O. acknowledges the funding provided by the Academy of Finland (MetaStem Center of Excellence grant 312437), the Novo Nordisk Foundation, the Sigrid Juselius Foundation and the Wellcome Collaborative Award in Science. H.I. acknowledges the funding provided by the Doctoral Programme in Biomedicine (DPBM), Orion Research Foundation, Paulon Säätiö Foundation, Diabetestutkimussäätiö Foundation, Biomedicum Helsinki Foundation, the Finnish Cultural Foundation and Magnus Ehrnrooth Foundation. The quality-check and library preparation of the RNAseq samples were provided by the Biomedicum Functional Genomics Unit at the Helsinki Institute of Life Science and Biocenter Finland at the University of Helsinki. Novaseq sequencing was performed by the Sequencing laboratory of Institute for Molecular Medicine Finland FIMM Technology Centre, University of Helsinki. A.T. acknowledges project grants from the Swedish Research Council, Novo Nordisk Foundation, Diabetes Wellness Sverige, the Swedish Diabetes Foundation, Barndiabetesfonden, Family Ernfors Foundation and the strategic grant consortium Excellence of Diabetes Research in Sweden (EXODIAB).

We are grateful for the family of the Mitchell-Riley Syndrome patient for participating and we want to acknowledge the participants and investigators of the FinnGen study. The FinnGen project is funded by two grants from Business Finland (HUS 4685/31/2016 and UH 4386/31/2016) and the following industry partners: AbbVie Inc., AstraZeneca UK Ltd, Biogen MA Inc., Bristol Myers Squibb (and Celgene Corporation & Celgene International II Sàrl), Genentech Inc., Merck Sharp & Dohme LCC, Pfizer Inc., GlaxoSmithKline Intellectual Property Development Ltd., Sanofi US Services Inc., Maze Therapeutics Inc., Janssen Biotech Inc, Novartis AG, and Boehringer Ingelheim International GmbH. Following biobanks are acknowledged for delivering biobank samples to FinnGen: Auria Biobank, THL Biobank, Helsinki Biobank, Biobank Borealis of Northern Finland, Finnish Clinical Biobank Tampere, Biobank of Eastern Finland, Central Finland Biobank, Finnish Red Cross Blood Service Biobank, Terveystalo Biobank and Arctic Biobank. All Finnish Biobanks are members of BBMRI infrastructure. Finnish Biobank Cooperative (FINBB) is the coordinator of BBMRI-ERIC operations in Finland. The Finnish biobank data can be accessed through the Fingenious® services managed by FINBB.

## Author contributions statement

HI conceived and conceptualized the study, generated the H1 hESCs lines, standardized the differentiation protocols, performed the experiments, analyzed data and wrote the original manuscript. DB and SE generated the iPSCs corrected cell-lines and DB participated in manuscript writing. JSV and HM performed and analyzed animal experiments. MOH, OD and PL performed and analyzed cell electrophysiology experiments. JK analyzed the bulk RNAseq datasets. OPD analyzed the FinnGen data and TT supervised the analysis. VL conducted the morphometric analyses pipelines. TB and VC participated in manuscript writing. JU performed the insulin and C-peptide ELISA measurements. SE participated in generating the genetically-edited clones, RNA purification and RT-qPCR experiments. PJM is the pediatric endocrinologist in charge of the patient case. SB and AT supervised the cell electrophysiology experiments, and participated in manuscript writing. TO conceived and supervised the study, provided resources, acquired funding and participated in manuscript writing.

