## Supplementary data for "*RFX6* haploinsufficiency predisposes to diabetes through impaired beta cell functionality"

Hazem Ibrahim *et al.*

**This PDF file includes:**

Supplementary Figs. 1 to 9
Supplementary Tables 1 to 6

Supplementary Fig. 1

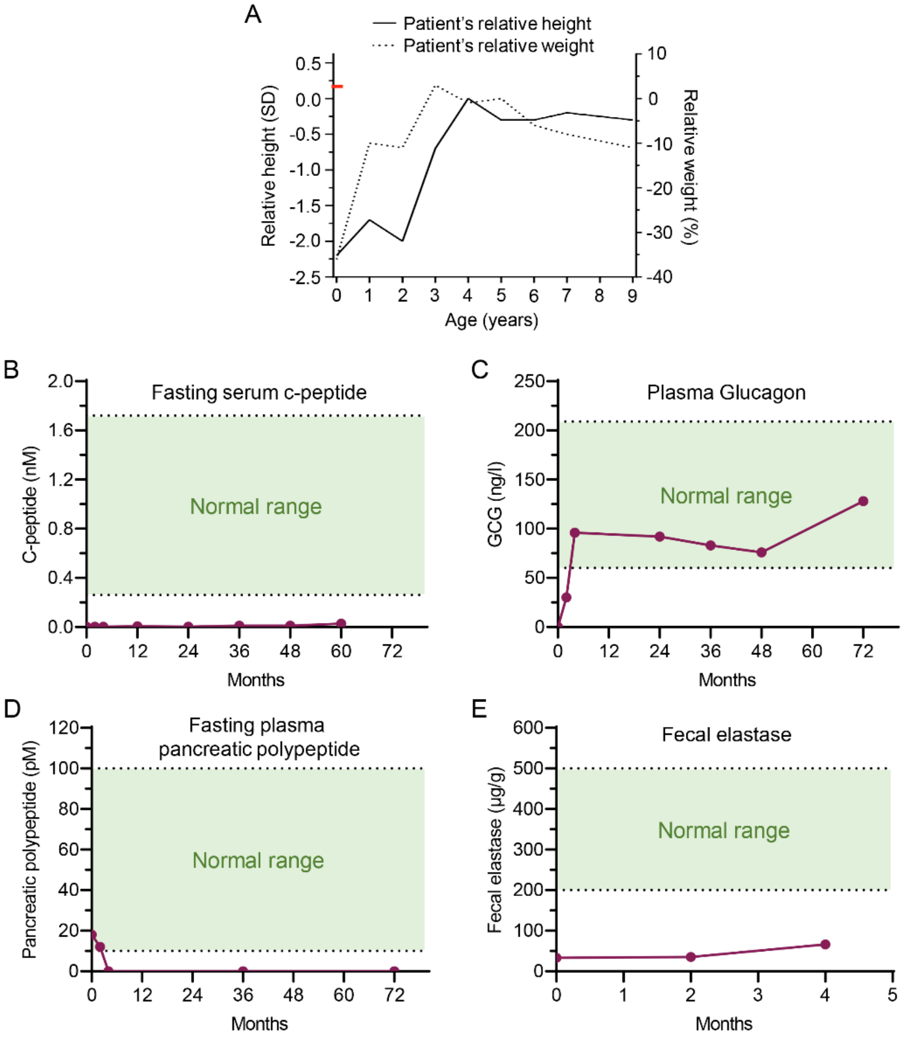


**Supplementary Fig. 1 Clinical manifestations of the homozygous *RFX6* patient.**

(**A**) Growth chart showing the patient's relative height and weight. Patient was born at gestation week 38+1, height was 47 cm and weight was 1800 g. Parental target relative height (+0.3 SD; red mark on left y axis). Relative height is depicted with solid line (left y axis) and relative weight (i.e. weight-for height) with dotted line (right y axis).
(**B**) Fasting serum c-peptide levels in nM during the first 5 years of age (normal range 0.26-1.72 nM).
(**C**) Plasma Glucagon levels in ng/l during the first 6 years of age (normal range 60-209 ng/l).
(**D**) Fasting plasma pancreatic polypeptide levels in pM during the first 6 years of age (normal range 10-100 pM).
(**E**) Fecal elastase levels in µg/g for the first 4 months after birth (normal level >200 µg/g).

Supplementary Fig. 2

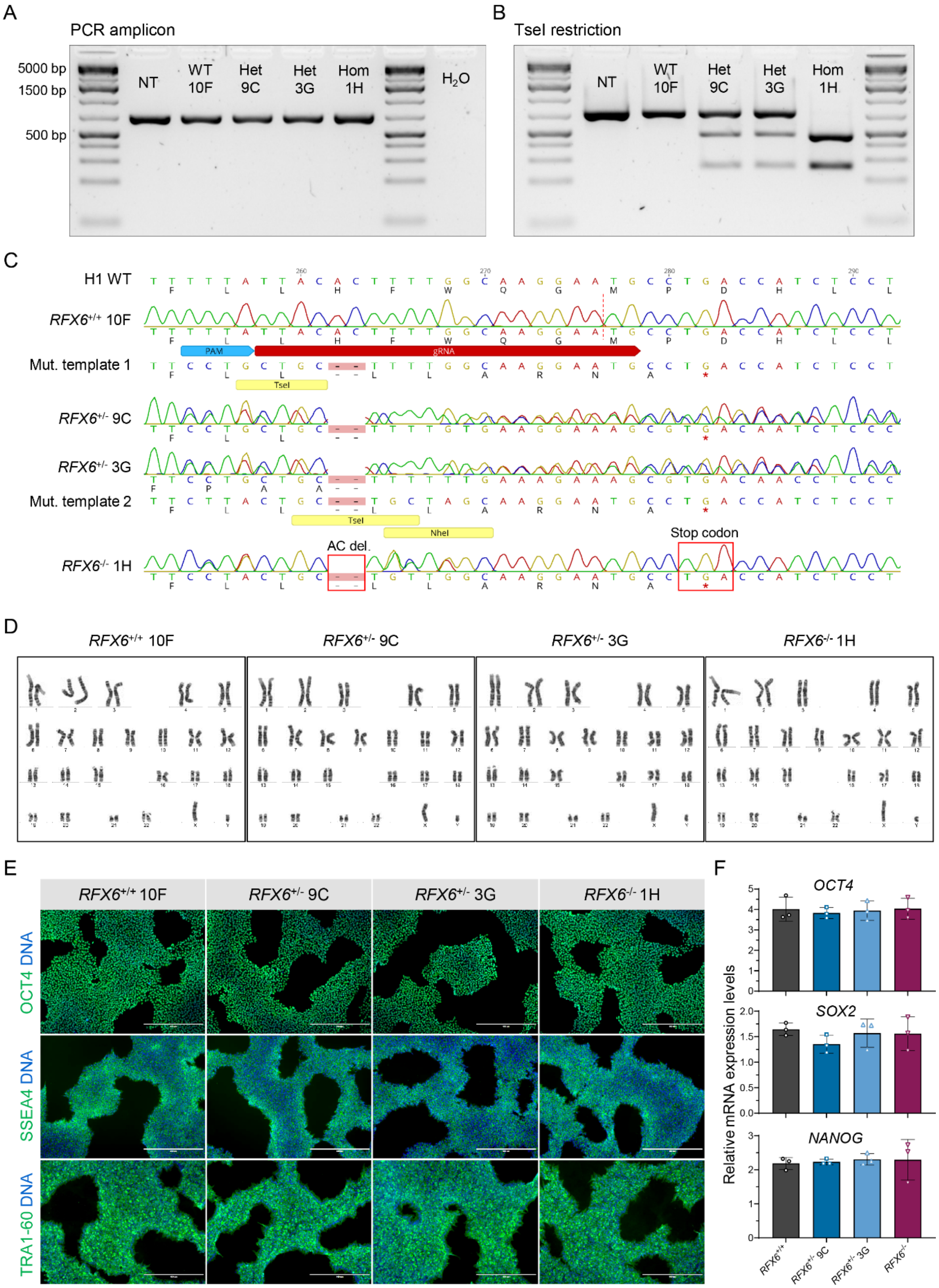

**Supplementary Fig. 2 *RFX6* patient PTV introduction in H1 hESCs using CRISPR-Cpf1.**

(**A**) PCR gel electrophoresis showing 729 bp amplicons in all the cell-lines.
(**B**) PCR gel electrophoresis of the TseI restricted amplicons showing the restricted bands of the edited alleles 264 + 465 bp in heterozygous and homozygous cell-lines.
(**C**) Sanger sequencing showing the PTV introduction heterozygously and homozygously using CRISPR-Cpf1 gRNAs and mutation templates.
(**D**) Normal karyotype visualized with G-banding for the four generated clones.
(**E**) Immunocytochemistry for pluripotency factors OCT4, TRA 1-60 and SSEA4 for the four clones, scale bars, 400 µm.
(**F**) Relative gene expression levels of pluripotency factors OCT4, SOX2 and NANOG for the four clones.
Statistical significance was measured using one-way ANOVA with Tukey's test for multiple comparisons correction in (F). Error bars represent ±SD from the mean.

#### Supplementary Fig. 3

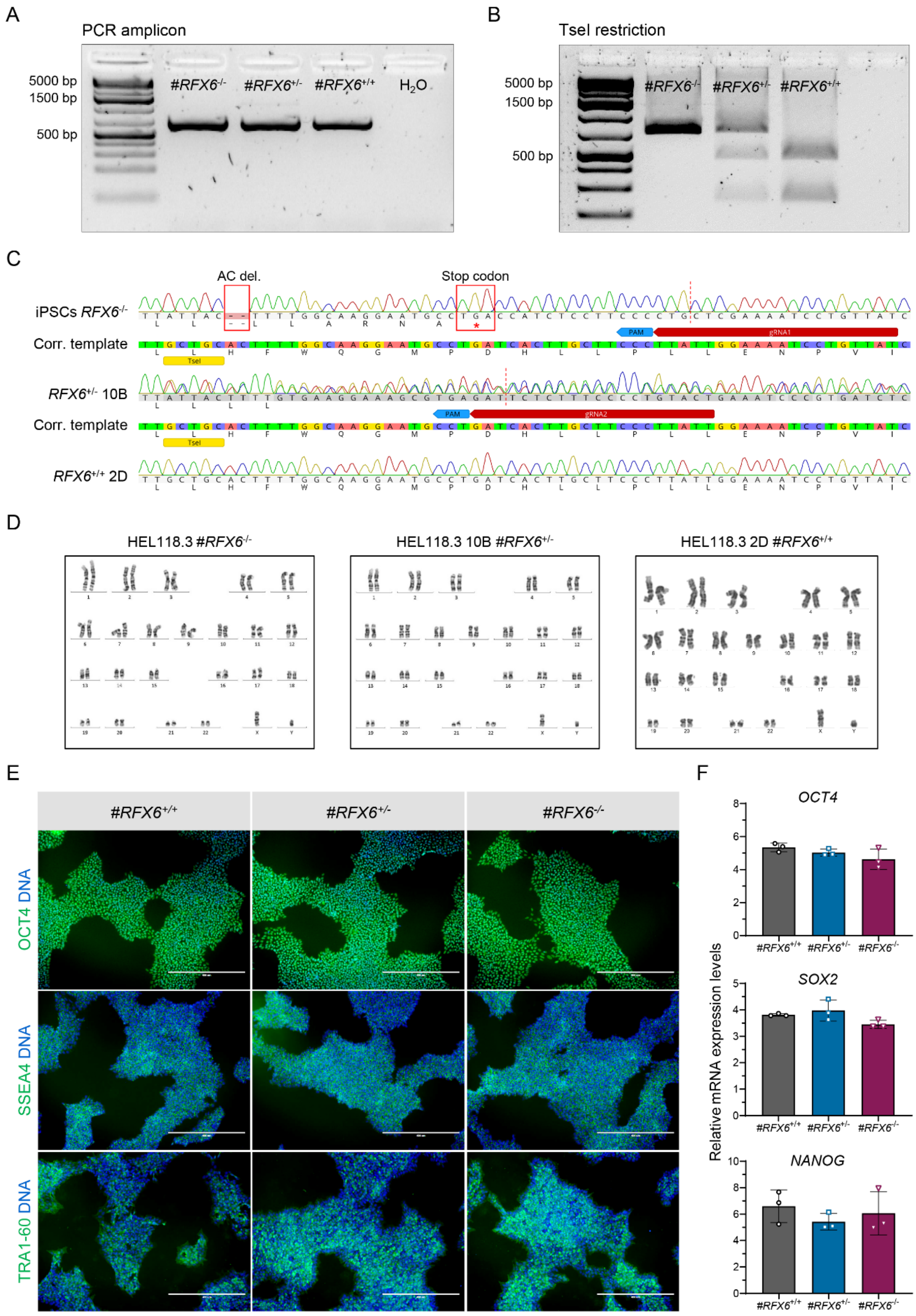

**Supplementary Fig. 3 *RFX6* PTV correction in patient derived iPSCs using CRISPR-Cas9.**

(**A**) PCR gel electrophoresis showing 729 bp amplicons of *RFX6* in all the cell-lines.
(**B**) PCR gel electrophoresis of the TseI restricted amplicons showing the restricted bands of the corrected alleles 264 + 465 bp in heterozygous and homozygous corrected cell-lines.
(**C**) Sanger sequencing showing the PTV correction of the patient derived iPSCs using CRISPR-Cas9 gRNAs and a correction template.
(**D**) Normal karyotype visualized with G-banding for the three clones.
(**E**) Immunocytochemistry for pluripotency factors OCT4, TRA 1-60 and SSEA4 for the three clones, scale bars, 400 µm.
(**F**) Relative gene expression levels of pluripotency factors OCT4, SOX2 and NANOG for the three clones (n = 3).
Statistical significance was measured using one-way ANOVA with Tukey's test for multiple comparisons correction in (F). Error bars represent ±SD from the mean.

Supplementary Fig. 4

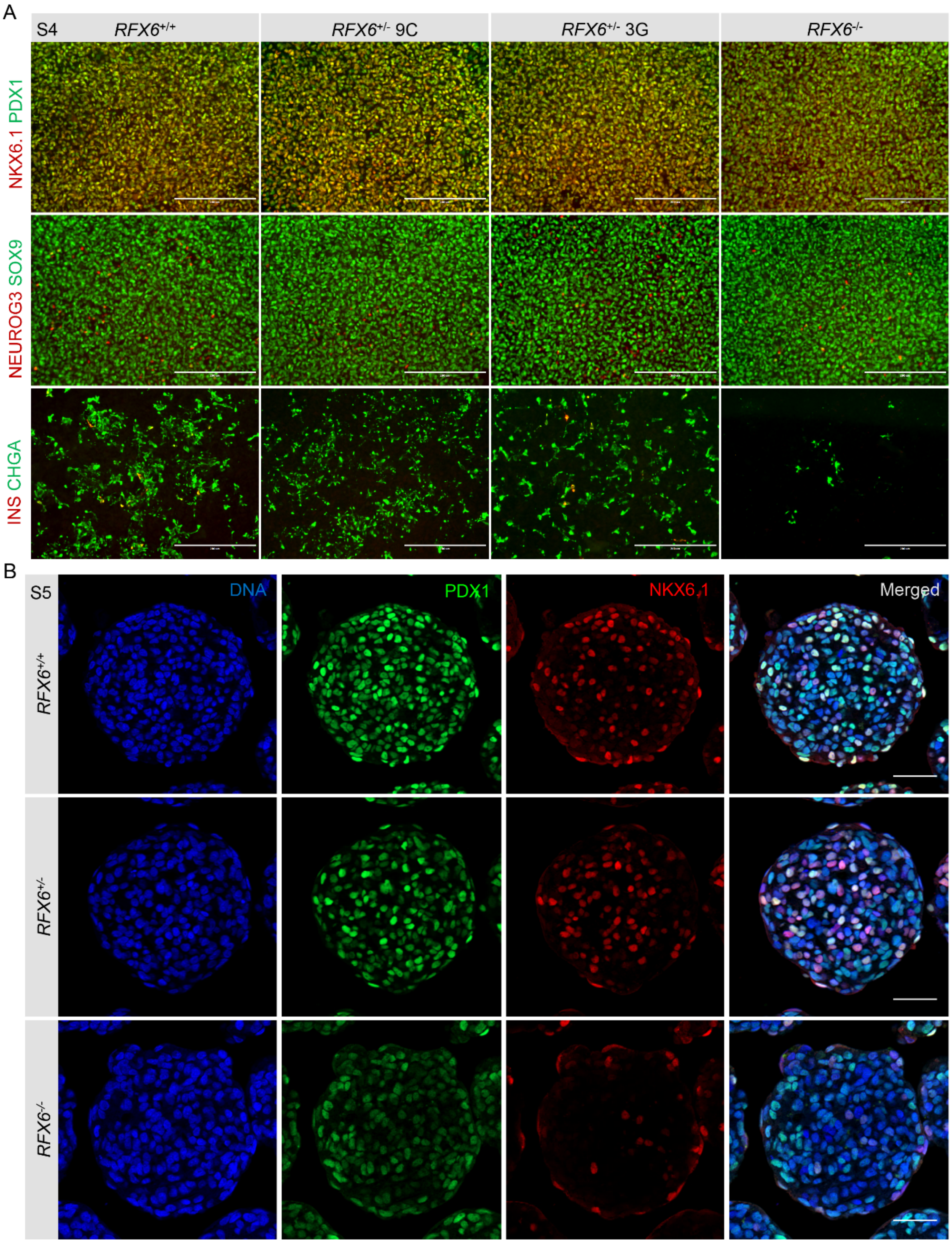


**Supplementary Fig. 4 Significant reduction in pancreatic endocrine precursors in homozygous *RFX6* PTV.**

(**A**) Immunocytochemistry showing PDX1$^+$ NKX6.1$^+$, SOX9$^+$ NEUROG3$^+$ and CHGA$^+$ INS$^+$ cells at S4, scale bars, 200 µm.
(**B**) Immunohistochemistry showing SOX9$^+$ and NEUROG3$^+$ cells at S5, scale bars, 50 µm.

Supplementary Fig. 5

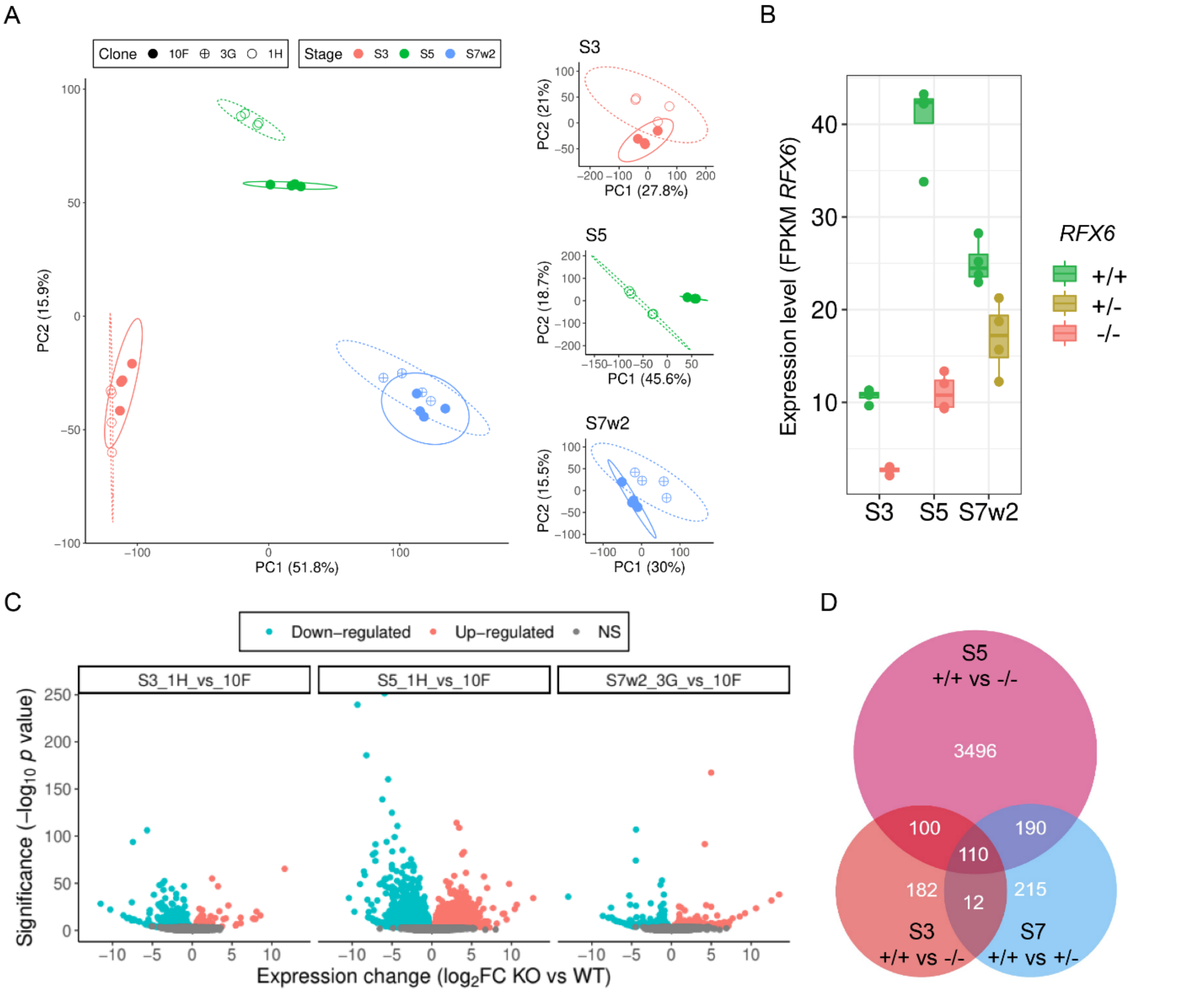


**Supplementary Fig. 5 RNAseq analysis of H1 *RFX6* differentiated cells.**

(**A**) Principal component (PC) analysis of the bulk RNA-seq samples. Filled circles, *RFX6*$^{+/+}$ 10F cells; crossed circles, *RFX6*$^{+/-}$ 3G cells; empty circles, *RFX6*$^{-/-}$ 1H cells (n=4).

(**B**) FPKM values for *RFX6* expression at different differentiation stages between the different genotypes (n=4).

(**C**) Volcano plot for differentially expressed genes for *RFX6*$^{+/+}$ and *RFX6*$^{-/-}$ at S3 and S5, and for *RFX6*$^{+/+}$ and *RFX6*$^{+/-}$ at S7w2. Significantly downregulated genes, iris blue; upregulated genes, soft red.

(**D**) Venn diagram showing common or unique differentially expressed genes across the genotypes.

### Supplementary Fig. 6

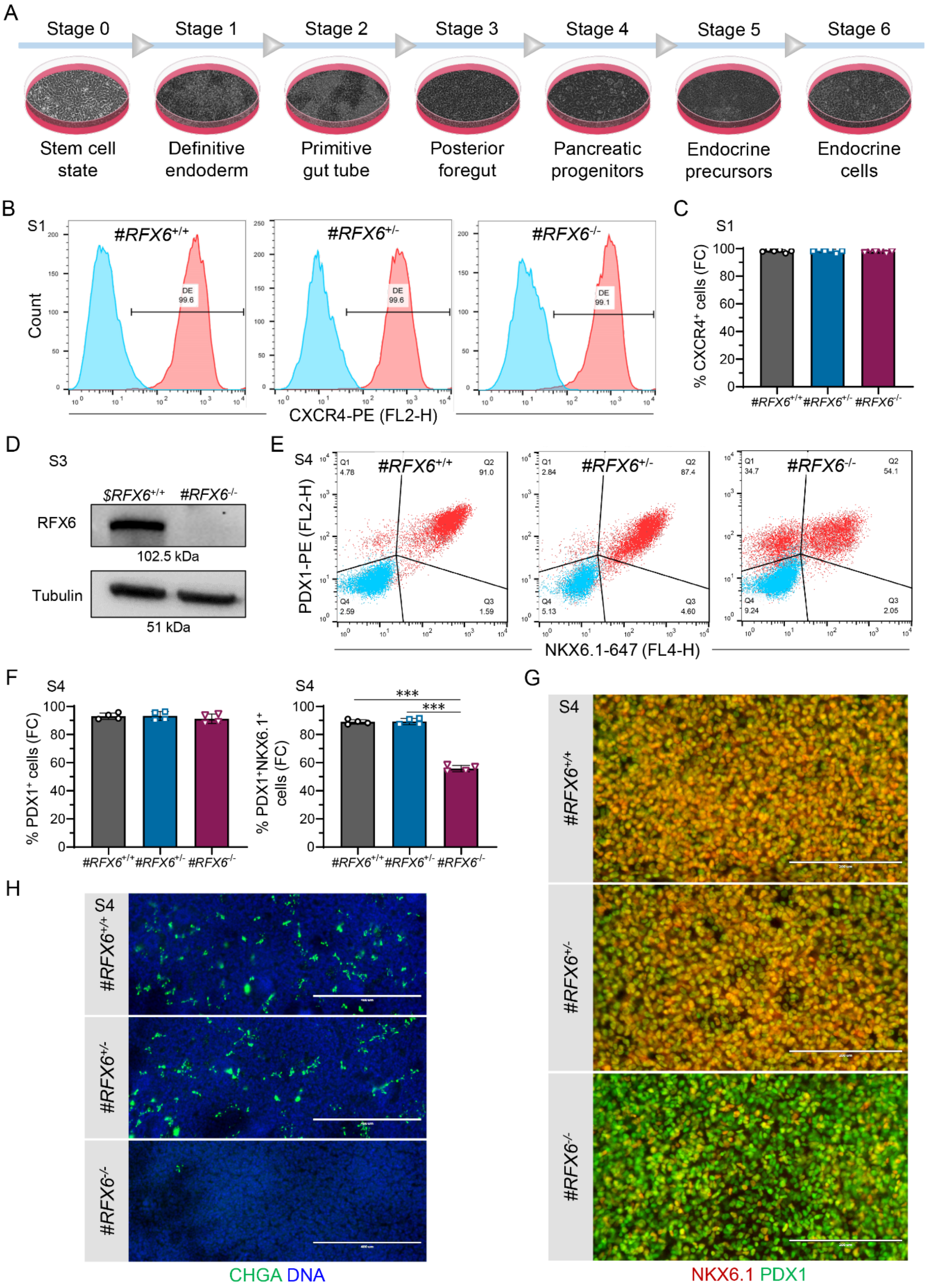

**Supplementary Fig. 6 Reduced numbers of pancreatic progenitor cells in patient-derived iPSCs.**

(**A**) Schematic of pancreatic endocrine differentiation protocol for 6 stages in monolayer.
(**B**) Flow cytometry analysis for the expression of $CXCR4^+$ definitive endoderm cells at the end of S1.
(**C**) Quantification of (B) (n = 6).
(**D**) Immunoblot for RFX6 and β-Actin at S3 of WT iPSCs Hel 46.11 (*$RFX6*$^{+/+}$) compared to the patient cell-line (*#RFX6*$^{-/-}$).
(**E**) Flow cytometry analysis for $PDX1^+$ and $NKX6.1^+$ cells at the end of S4.
(**F**) Quantification of (E) (n = 4).
(**G**) Immunocytochemistry for $PDX1^+$ and $NKX6.1^+$ at the end of S4, scale bars, 200 µm.
(**H**) Immunocytochemistry for $CHGA^+$ at the end of S4, scale bars, 400 µm.
Statistical significance was measured using one-way ANOVA with Tukey's test for multiple comparisons correction in (C and F). Error bars represent ±SD from the mean, *p<0.05, **p<0.01, ***p<0.001.

Supplementary Fig. 7

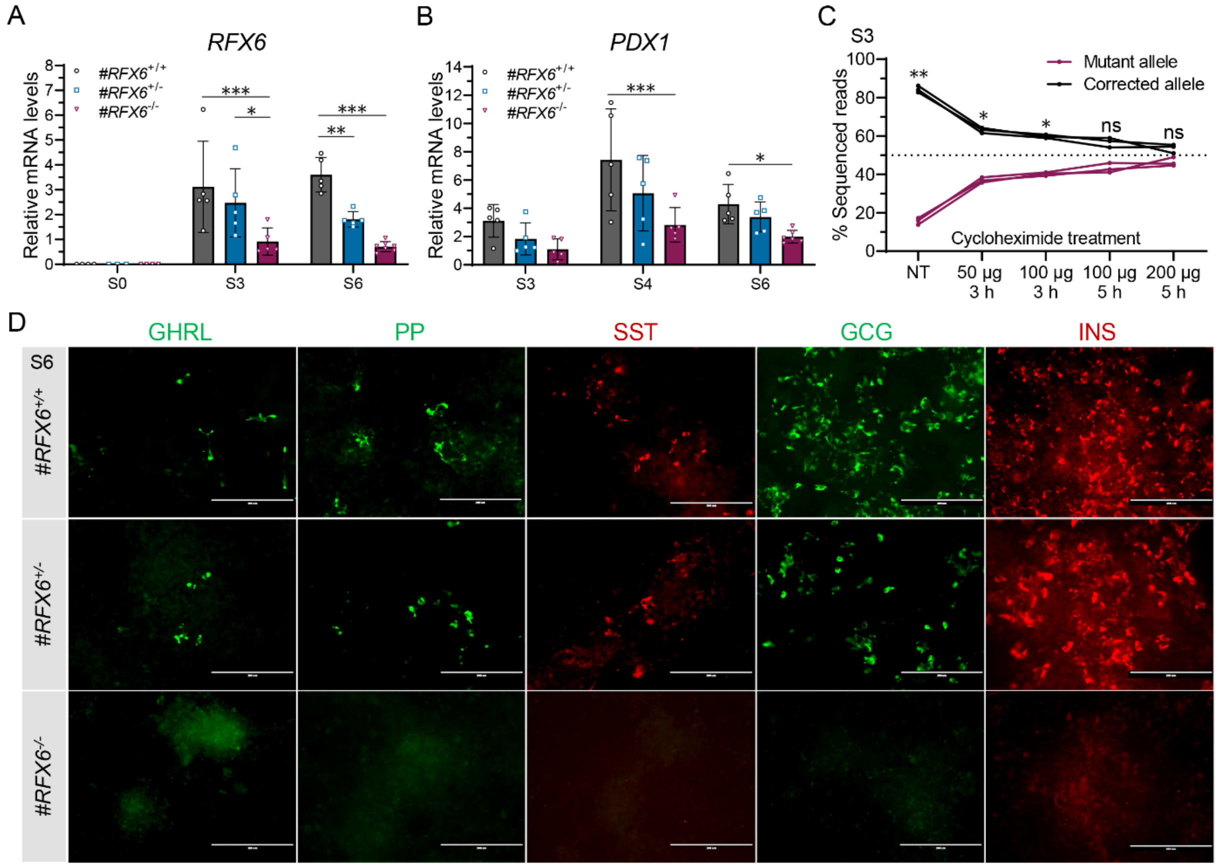


**Supplementary Fig. 7 *RFX6* PTV nonsense-mediated mRNA decay and abolished production of pancreatic endocrine cells from patient-derived iPSCs.**

(**A**) Relative gene expression levels of *RFX6* for the three isogenic cell-lines at S0, S3 and S6 (n=3–7).
(**B**) Relative gene expression levels of *PDX1* for the three isogenic cell-lines at S3, S4 and S6 (n=5–6).
(**C**) Percentage of *RFX6* cDNA reads of the variant and corrected alleles at S3 of *RFX6*$^{+/-}$ cell-line treated with different doses of the NMD inhibitor cycloheximide (n=3).
(**D**) Immunocytochemistry at S6 for the five pancreatic endocrine cell types. Ghrelin (GHRL), Pancreatic polypeptide (PP), Somatostatin (SST), Glucagon (GCG) and Insulin (INS), scale bars, 200 µm.
Statistical significance was measured using two-way ANOVA with Tukey's test for multiple comparisons correction in (A and B), and using paired parametric multiple t-tests in (C). Error bars represent ±SD from the mean, *p<0.05, **p<0.01, ***p<0.001.

Supplementary Fig. 8

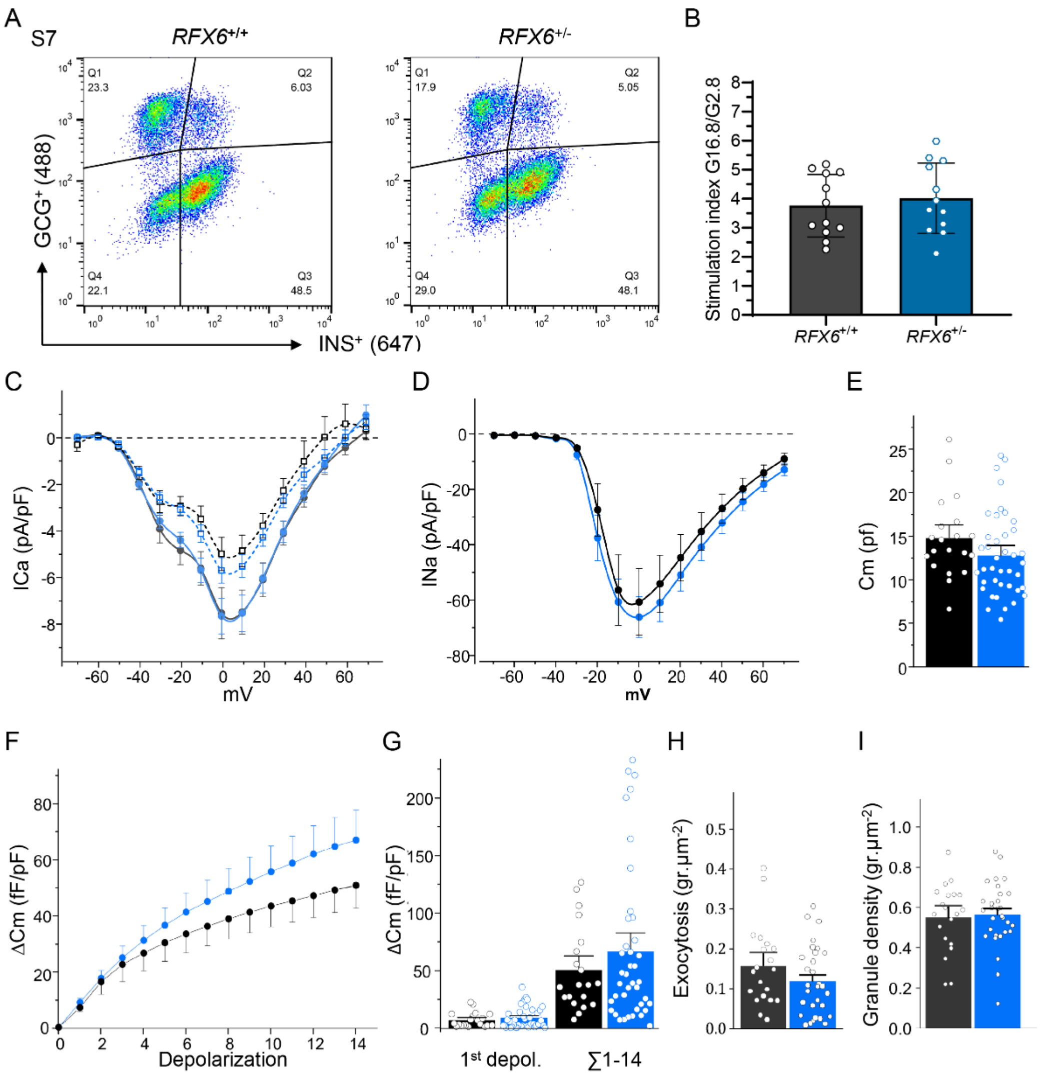


**Supplementary Fig. 8 Voltage-dependence and exocytosis of RFX6 SC-islets.**

(**A**) Flow cytometry analysis for $GCG^+$ cells (y-axis) and $INS^+$ cells (x-axis) at S7w2.
(**B**) Insulin secretion stimulation index from static GSIS assays (G16.8 mM/G2.8 mM) (n=12).
(**C**) Voltage-dependence of $Ca^{2+}$-currents in wt (black, n=20 cells) and PTV (blue, n=37 cells) *RFX6*$^{+/-}$ beta-cells, in absence (solid lines) or presence of 20 µM nifedipine.
(**D**) Voltage-dependence of $Na^+$-currents in the same cells as in C.
(**E**) Initial whole cell capacitance, as measure of cell size.
(**F**) Exocytosis stimulated by a train of voltage-clamp depolarizations, measured as capacitance increase in wt (black) and PTV (blue, n=40 cells) *RFX6*$^{+/-}$ beta-cells.
(**G**) Quantfication of capacitance changes in F during the first (left) and the sum of all depolarizations.
(**H**-**I**) Exocytosis (**H**) and initial docked granules (**I**) measured by TIRF microscopy of NPY-EGFP labeled insulin granules in wt (black, n=20 cells) and PTV (blue, n=30 cells) *RFX6*$^{+/-}$ beta-cells. Cells were stimulated by elevating $K^+$ for 40s; data are normalized to cell footprint area.

Supplementary Fig. 9

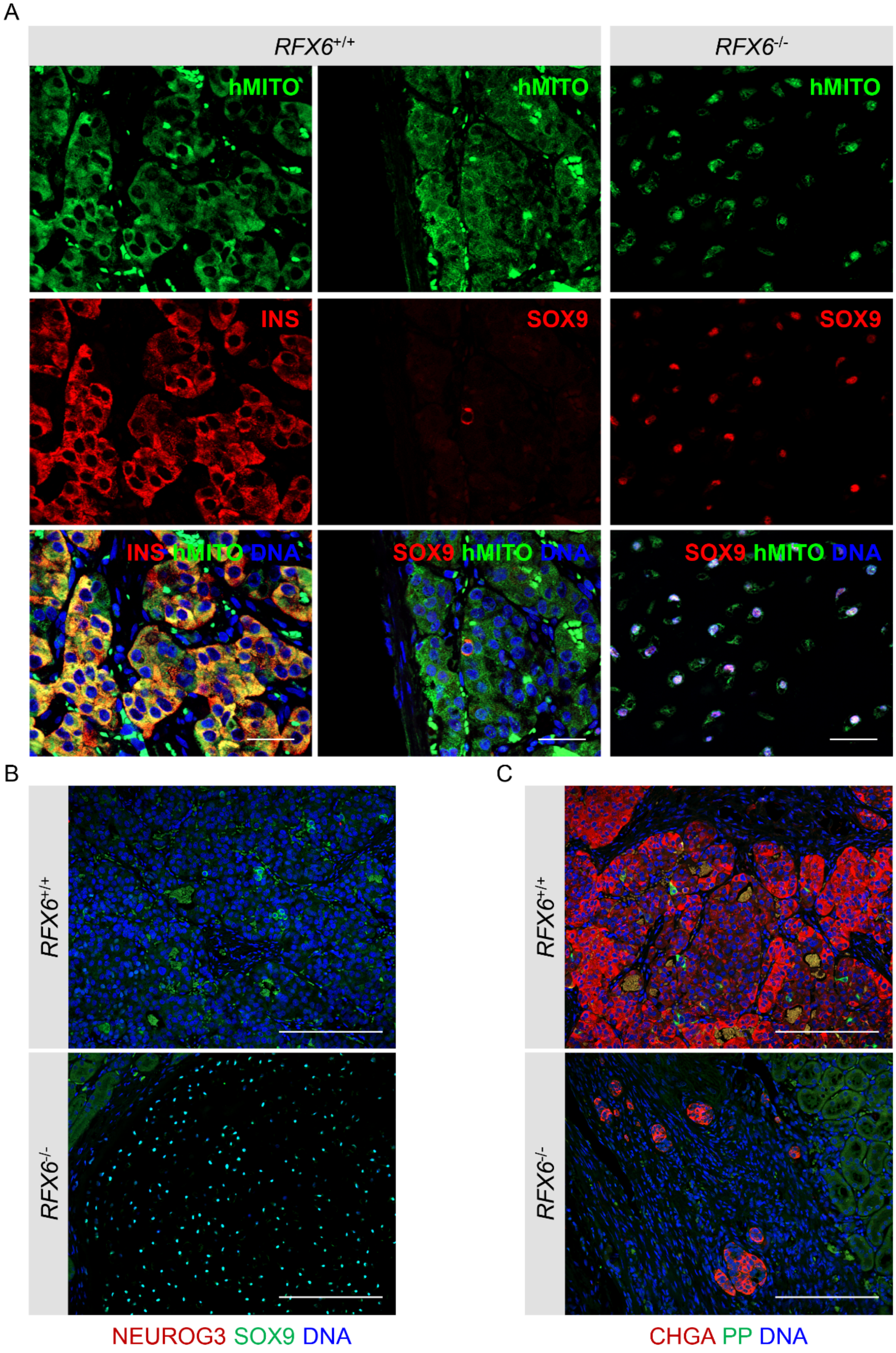

**Supplementary Fig. 9 Sustained expression of SOX9 but not NEUROG3 in homozygous *RFX6*$^{-/-}$ cells post implantation.**

(**A**) Immunohistochemistry showing human specific mitochondria (hMITO$^{+}$), INS$^{+}$ and SOX9$^{+}$ cells at month 3 post implantation, scale bars, 50 µm.
(**B**) Immunohistochemistry showing SOX9$^{+}$ and NEUROG3$^{+}$ cells at month 3 post implantation, scale bars, 200 µm.
(**C**) Immunohistochemistry showing PP$^{+}$ and CHGA$^{+}$ cells at month 3 post implantation, scale bars, 200 µm.

**Supplementary Table 1 Primers for PCR, CRISPR off-targets and NGS sequencing**

| Primer | Sequence |
|---|---|
| RFX6 Fw | GGCAGTGTTAAAAGGATGCTTTGG |
| RFX6 Rv | GGGTGCCTAATTCTGCTTTCTG |
| RFX6 dsDNA Mutation TseI 200bp Fw | TATTATATATACTTTCATGTTCTGTTCTTAATGAGTTCATGTTAAAGAAAAAAATCTTAACATACTTTTTACTCTGCAACTAGATCCAGCATTTTTTGCTGCTTTTGG |
| RFX6 dsDNA Mutation TseI 200bp Rv | TACCTTATAAAGAATTGAGTCACAAACACAGAAAATATCAATGATAACAGGATTTTCCAATAAGGGAAGCAAGTGATCAGGCATTCCTTGCCAAAAGCAGCAAAAAAT |
| RFX6 Cpf1 Off-target1 Fw | TCCCAGGATTAAGTGTGGGG |
| RFX6 Cpf1 Off-target1 Rv | TCAATATTTGGTACCTGCCAACT |
| RFX6 Cpf1 Off-target2 Fw | TCCTGCACAGAGGTCCTACA |
| RFX6 Cpf1 Off-target2 Rv | AGCATCATGGGTGTTCGACT |
| RFX6 Cpf1 Off-target3 Fw | CAGGGAACGAGGGAGAACTG |
| RFX6 Cpf1 Off-target3 Rv | GAGTTTCCTAGGTGGGTGCC |
| RFX6 cDNA NGSseq Fw | *ACACTCTTTCCCTACACGACGCTCTTCCGATCT*ACTCTGCAACTAGATCCAGCATTTT |
| RFX6 cDNA NGSseq Rv | *AGACGTGTGCTCTTCCGATCT*AAGAATTGAGTCACAAACACAGAAA |

**Supplementary Table 2 Differentiation protocol and media formulations for hESCs**

<table>
<tr><th>Stage</th><th>Reagent</th><th>Medium</th></tr>
<tr><td>Stage 1<br>3 days</td><td>d0: 100 ng/ml human Activin A, 3 μM CHIR<br>d1: 100 ng/ml human Activin A, 0.3 μM CHIR<br>d2: 100 ng/ml human Activin A</td><td rowspan="2">Basal 1: MCDB131 (10372-019, Life Technologies), 2 mM Glutamax, 1.5 g/L NaHCO3 (Sigma-Aldrich), 0.5% BSA fraction V Fatty acid free (Sigma-Aldrich), 10 mM glucose</td></tr>
<tr><td>Stage 2<br>3 days</td><td>0.25 mM Ascorbic acid, 50 ng/ml FGF7</td></tr>
<tr><td>Stage 3<br>2 days</td><td>0.25 mM Ascorbic acid, 50 ng/ml FGF7, 0.25 μM SANT1, 1 μM Retinoic Acid, 100 nM LDN, 200 nM and TPB</td><td rowspan="2">Basal 2: MCDB131, 2 mM Glutamax, 2.5 g/L NaHCO3, 2% BSA fV, 10 mM glucose, 1:200 ITS-X (51500-056, Life Technologies)</td></tr>
<tr><td>Stage 4<br>4 days</td><td>0.25 mM Ascorbic acid, 2 ng/ml FGF7, 0.25 μM SANT1, 0.1 μM Retinoic Acid, 200 nM LDN, 100 ng/ml EGF, 10 mM Nicotinamide and 10 μM ROCKi</td></tr>
<tr><td>Stage 5<br>4 days</td><td>0.25 μM SANT1, 0.05 μM Retinoic acid, 100 nM LDN, 10 μM ALK5inhII, 1 μM GC1, 20 ng/mL Betacellulin, 100 nM GSiXX and 5 μM ROCKi</td><td rowspan="2">Basal 3: MCDB131, 2 mM Glutamax, 1.5 g/L NaHCO3, 2% BSA fV, 20 mM glucose, 1:200 ITS-X, 10 μg/mL Heparin (H3149, Sigma-Aldrich), 10 μM Zinc Sulfate</td></tr>
<tr><td>Stage 6<br>8 days</td><td>100 nM LDN, 10 μM ALK5inhII, 1 μM GC1 and 100 nM GSiXX</td></tr>
<tr><td>Stage 7<br>21 days</td><td>1 mM N-Acetylcysteine, 10 nM Tri-iodothyronine (T3) and 0.5 μM Aurora kinase inhibitor ZM447439</td><td>CMRL-modified: CMRL 1066 (15-110-CVR, Corning), 2% BSA fV, 2 mM Glutamax, 1:200 ITS-X, Heparin 10 μg/ml, 10 μM Zinc Sulfate, 5 mM Sodium Pyruvate (Lonza), 1:2000 chemically defined lipid concentrate, 1:2000 medium trace elements A, 1:2000 medium trace elements B</td></tr>
</table>

**Supplementary Table 3 Differentiation protocol and media formulations for iPSCs differentiation**

| Stage | Reagent | Medium |
|---|---|---|
| Stage 1<br>2 days | d0: 100 ng/ml human Activin A, 3 μM CHIR<br>d1: 100 ng/ml human Activin A, 0.3 μM CHIR | Basal 1: MCDB131 (10372-019, Life Technologies), 2 mM Glutamax, 1.5 g/L NaHCO3 (Sigma-Aldrich), 0.5% BSA fraction V Fatty acid free (Sigma-Aldrich), 10 mM glucose |
| Stage 2<br>3 days | 0.25 mM Ascorbic acid, 50 ng/ml FGF7 | |
| Stage 3<br>2 days | 0.25 mM Ascorbic acid, 50 ng/ml FGF7, 0.25 μM SANT1, 1 μM Retinoic Acid, 100 nM LDN, 200 nM and TPB | Basal 2: MCDB131, 2 mM Glutamax, 2.5 g/L NaHCO3, 2% BSA fV, 10 mM glucose, 1:200 ITS-X (51500-056, Life Technologies) |
| Stage 4<br>4 days | 0.25 mM Ascorbic acid, 2 ng/ml FGF7, 0.25 μM SANT1, 0.1 μM Retinoic Acid, 200 nM LDN, 100 ng/ml EGF and 10 mM Nicotinamide | |
| Stage 5<br>4 days | 0.25 μM SANT1, 0.05 μM Retinoic acid, 100 nM LDN, 10 μM ALK5inhII, 1 μM GC1, 20 ng/mL Betacellulin and 100 nM GSiXX | Basal 3: MCDB131, 2 mM Glutamax, 1.5 g/L NaHCO3, 2% BSA fV, 20 mM glucose, 1:200 ITS-X, 10 μg/mL Heparin (H3149, Sigma-Aldrich), 10 μM Zinc Sulfate |
| Stage 6<br>12 days | 100 nM LDN, 10 μM ALK5inhII, 1 μM GC1, 100 nM GSiXX and 10 μM ROCKi | |

**Supplementary Table 4 Differentiation growth factors and reagents**

| Chemical/Reagent | Company name | Catalog# |
|---|---|---|
| ROCKi Y-27632 | Selleckchem | Cat# S1049 |
| Activin A | QKine Ltd, Cambridge, UK | Cat# Qk001 |
| CHIR | Tocris | Cat# 4423 |
| Ascorbic acid | Sigma-Aldrich | Cat# A4544 |
| FGF7 | Genscript | Cat# Z03047 |
| SANT1 | Sigma-Aldrich | Cat# 4572 |
| LDN | Selleckchem | Cat# S2618 |
| Retinoic acid | Sigma-Aldrich | Cat# R2625 |
| TPB | Santa Cruz | Cat# sc-204424 |
| EGF | Peprotech | Cat# AF-100-15 |
| Nicotinamide | Sigma-Aldrich | Cat# N0636 |
| ALK5inhII | Selleckchem | Cat# S7233 |
| GC1 | Tocris | Cat# 4554 |
| GSiXX | Millipore | Cat# 56578 |
| N-Acetylcysteine | Sigma-Aldrich | Cat# A9165 |
| Tri-iodothyronine (T3) | Sigma-Aldrich | Cat# T6397 |
| Aurora kinase inhibitor ZM447439 | Selleckchem | Cat# S1103 |
| Glutamax | Life Technologies | Cat# 35050038 |
| Heparin | Sigma-Aldrich | Cat# H3149 |
| Zinc Sulfate | Sigma-Aldrich | Cat# Z0251 |
| Chemically defined lipid concentrate | Invitrogen | Cat# 11905-031 |
| Medium trace elements A | Cellgro | Cat# 25-021-CI |
| Medium trace elements B | Cellgro | Cat# 99-175-CI |
| Cycloheximide | Sigma | Cat# 01810-1G |

**Supplementary Table 5 Antibodies used for immunofluorescence, flow cytometry and immunoblotting**

| Antibody | Company | Catalog# | Dilution |
|---|---|---|---|
| Rabbit anti-OCT4 Rabbit | Santa Cruz Biotechnology | Cat# sc-9081; RRID: AB_2167703 | 1:250 ICC |
| Rabbit anti-SSEA4 mAb DyLight™ 488 | ThermoFisher Scientific | Cat# MA1-021-D488; RRID: AB_2536688 | 1:100 ICC |
| Mouse anti-TRA 1-60 mAb | ThermoFisher Scientific | Cat# MA1-023; RRID: AB_2536699 | 1:100 ICC |
| Goat anti-PDX1 | R&D systems | Cat# AF2419; RRID: AB_355257 | 1:500 IHC |
| Mouse anti-NKX6.1 | DSHB Hybridoma | Cat# F55A10; RRID: AB_532378 | 1:250 IHC |
| Sheep anti Human Neurogenin-3 | R&D systems | Cat# AF3444; RRID: AB_2149527 | 1:250 ICC, IHC |
| Rabbit anti-SOX9 | Sigma-Aldrich | Cat# AB5535; RRID: AB_2239761 | 1:500 IHC |
| Mouse anti-GCG antibody | Sigma-Aldrich | Cat# G2654; RRID: AB_259852 | FC-1:160 IHC- 1:500 |
| Guinea pig anti-Insulin | Dako | Cat# A0564; RRID: AB_10013624 | 1:1000 IHC |
| Rabbit anti-Somatostatin | Dako | Cat# A0566; RRID: AB_2688022 | 1:500 ICC |
| Goat anti-Pancreatic polypeptide | Sigma-Aldrich | Cat# SAB2500747; RRID: AB_10611538 | 1:500 |
| Goat anti-Ghrelin | Santa Cruz | Cat# sc-10368; RRID: AB_2232479 | 1:500 ICC |
| Mouse anti-CHGA | Dako | Cat# M0869; RRID: AB_2081135 | FC-1:160 IHC- 1:500 |
| Rabbit anti-SLC18A1 | Sigma-Aldrich | Cat# HPA063797; RRID: AB_2685125 | 1:250 IHC |
| Mouse anti-human Mitochondria clone 113-1 | Sigma-Aldrich | Cat# MAB1273; RRID: AB_94052 | 1:150 IHC |
| Mouse Anti-CD184 (CXCR4) Monoclonal Antibody | BD Biosciences | Cat# 555974; RRID: AB_396267 | 1:10 FC |
| Mouse IgG2a, kappa Isotype Control, Phycoerythrin Conjugated | BD Biosciences | Cat# 5555749; RRID: AB_396091 | 1:10 FC |
| PE-Mouse anti PDX1 | BD Biosciences | Cat# 562161; RRID: AB_10893589 | 1:80 FC |
| Alexa Fluor 647 Mouse anti NKX6-1 | BD Biosciences | Cat# 563338; RRID: AB_2738144 | 1:80 FC |
| Alexa Fluor 647 Mouse IgG1 k isotype control | BD Biosciences | Cat# 557714; RRID: AB_396823 | 1:80 FC |
| Alexa Fluor 647 Rabbit anti Insulin (C27C9) | Cell Signaling Technology | Cat# 9008; RRID: AB_2687822 | 1:80 FC |

| | | | |
|---|---|---|---|
| Alexa Fluor 647 Rabbit IgG Isotype Control | Cell Signaling Technology | Cat# 3452S; RRID: AB_10695811 | 1:80 FC |
| Alexa FluoR 488 Donkey anti-Mouse IgG secondary antibody | ThermoFisher Scientific | Cat# A-21202; RRID: AB_141607 | 1:500 |
| Alexa FluoR 594 Goat anti-Guinea Pig IgG secondary antibody | ThermoFisher Scientific | Cat# A-11076; RRID: AB_2534120 | 1:500 |
| Alexa FluoR 594 Donkey anti-Sheep IgG secondary antibody | ThermoFisher Scientific | Cat# A- 11016; RRID: AB_2534083 | 1:500 |
| Alexa FluoR 488 Donkey anti-Rabbit IgG secondary antibody | ThermoFisher Scientific | Cat# A-21206; RRID: AB_2535792 | 1:500 |
| Alexa FluoR 488 Donkey anti-Mouse IgG secondary antibody | ThermoFisher Scientific | Cat# A-21203; RRID: AB_141633 | 1:500 |
| Alexa FluoR 488 Donkey anti-Goat IgG secondary antibody | ThermoFisher Scientific | Cat# A-11055; RRID: AB_2534102 | 1:500 |
| Mouse anti-alpha-Tubulin Mouse mAb | Sigma-Aldrich | Cat# T5168; (B-5-1-2); RRID: AB_477579 | 1:2000 WB |
| Mouse anti-β-actin-HRP mAb | Santa Cruz | Cat# sc-47778; (C4); RRID: AB_626632 | 1:1000 WB |
| Rabbit anti-RFX6 | Sigma-Aldrich | Cat# HPA037696; RRID: AB_2675616 | 1:100 WB |
| Anti-rabbit IgG, HRP-linked Antibody | Cell Signaling Technology | Cat# 7074; RRID: AB_2099233 | 1:2000 WB |
| Anti-mouse IgG, HRP-linked Antibody | Cell Signaling Technology | Cat# 7076; RRID: AB_330924 | 1:5000 WB |

**Supplementary Table 6 Primers for quantitative RT-PCR**

| Primer | NCBI Reference | Sequence |
|---|---|---|
| *PPIG* primer pair<br>Housekeeping gene | NM_004792 | Fw: TCTTGTCAATGGCCAACAGAG;<br>Rv: GCCCATCTAAATGAGGAGTTG; 84 bp |
| *OCT4* primer pair | NM_002701 | Fw: TTGGGCTCGAGAAGGATGTG;<br>Rv: TCCTCTCGTTGTGCATAGTCG; 91 bp |
| *SOX2* primer pair | NM_003106 | Fw: GCCCTGCAGTACAACTCCAT;<br>Rv: TGCCCTGCTGCGAGTAGGA; 85 bp |
| *NANOG* primer pair | NM_024865.2 | Fw: CTCAGCCTCCAGCAGATGC;<br>Rv: TAGATTTCATTCTCTGGTTCTGG; 94 bp |
| *RFX6* primer pair | NM_173560.1 | Fw: GCATGTCGAACTCCAGTCCT;<br>Rv: GGAGTCCAGGGTTTCTTGAG; 114 bp |
| *PDX1* primer pair | NM_000209.3 | Fw: AAGTCTACCAAAGCTCACGCG;<br>Rv: CGTAGGCGCCGCCTGC; 52 bp |
| *SOX9* primer pair | NM_000346 | Fw: ATCAAGACGGAGCAGCTGAG;<br>Rv: GGCTGTAGTGTGGGAGGTTG; 100 bp |
| *NEUROG3* primer pair | NM_020999 | Fw: GACGACGCGAAGCTCACCAA;<br>Rv: TACAAGCTGTGGTCCGCTAT; 98 bp |
